# High-throughput microcircuit analysis of individual human brains through next-generation multineuron patch-clamp

**DOI:** 10.1101/639328

**Authors:** Yangfan Peng, Franz X. Mittermaier, Henrike Planert, Ulf C. Schneider, Henrik Alle, Jörg R.P. Geiger

**Affiliations:** Institute of Neurophysiology, Charité – Universitätsmedizin Berlin, Germany; Department of Neurology, Charité – Universitätsmedizin Berlin, Germany; Department of Neurosurgery, Charité – Universitätsmedizin Berlin, Germany

## Abstract

Comparing neuronal microcircuits across different brain regions, species and individuals can reveal common and divergent principles of network computation. Simultaneous patch-clamp recordings from multiple neurons offer the highest temporal and subthreshold resolution to analyse local synaptic connectivity. However, its establishment is technically complex and the experimental performance is limited by high failure rates, long experimental times and small sample sizes. We introduce an in-vitro multipatch setup with an automated pipette pressure and cleaning system facilitating recordings of up to 10 neurons simultaneously and sequential patching of additional neurons. We present hardware and software solutions that increase the usability, speed and data throughput of multipatch experiments which allowed probing of 150 synaptic connections between 17 neurons in one human cortical slice and screening of over 600 connections in tissue from a single patient. This method will facilitate the systematic analysis of microcircuits and allow unprecedented comparisons at the level of individuals.

## Introduction

### Neuronal microcircuits represent the backbone for computations of the brain

While it is well established that information flow between the cortical layers is rather stereotypical (Douglas and Martin, 2004), the intralaminar connectivity between different classes of neurons is more complicated and heterogeneous across brain regions (Jiang et al., 2015; Peng et al., 2017; Song et al., 2005). Furthermore, it remains as yet unclear how these observed microcircuit topologies generalize across species, or whether and how inter-individual differences are represented at the level of neuronal microcircuits.

### Cortical resections from epilepsy or tumor patients represent a unique opportunity to analyse the human neuronal microcircuit

While specific cellular and synaptic properties in the human central nervous system have been described (Kalmbach et al., 2018; Molnár et al., 2016; Szegedi et al., 2016), distinctive features on the microcircuit level have yet to be resolved (Seeman et al., 2018). Availability of human brain tissue is scarce making it, at present, necessary to pool data across patients. This is problematic due to the high diversity in humans, in contrast to laboratory animals. Therefore, it is essential that we develop more efficient ways to generate large sample sizes from single patients. Apart from that, in non-human studies, increasing the data yield from individual animals would also efficiently promote the ethical principle to replace, reduce and refine animal experiments (Russell et al., 1959).

### To dissect microcircuits, multiple whole-cell patch-clamp (multipatch) recordings still represent the gold standard method compared to various other approaches

(Hochbaum et al., 2014; Packer et al., 2014). This method can reliably detect unitary excitatory and inhibitory synaptic connections due to its sub-millisecond and subthreshold resolution (Geiger et al., 1997). In addition, it allows for detailed electrophysiological and morphological characterization of recorded neurons and can be used across species without the need for genetic modifications in contrast to optogenetics or calcium imaging (Markram et al., 2004). These conditions enable the investigation of connection probabilities, amplitude distributions and higher order network statistics incorporating distance-dependencies (Song et al., 2005; Thomson and Lamy, 2007). Increasing the number of simultaneously recorded neurons can increase the number of probed synaptic connections considerably, generating larger sample sizes from fewer experiments.

### However, multipatch setups increase the complexity and time of experiments, necessitating automatisation

Various groups were able to increase the number of simultaneous recordings (Guzman et al., 2016; Jiang et al., 2015; Peng et al., 2017; Perin et al., 2011; Perin and Markram, 2013; Wang et al., 2015)), but the operation of multiple manipulators is challenging. To address this, several approaches have been reported that automate the patch-clamp process, utilizing automated pressure control systems and algorithms for manipulator movements guided by visual or electrical signals (Kodandaramaiah et al., 2018, 2016, 2012; Perin and Markram, 2013; Wu et al., 2016). While simultaneous recordings of up to 4 neurons are becoming increasingly popular, the use of more than 8 manipulators on a setup is currently limited to only a few labs, missing out on the opportunity to maximize the potential of this method.

### Cleaning pipettes for immediate reuse increases the size of recorded clusters of neurons and enables a more complete view of the microcircuit

The maximum number of simultaneously recorded neurons is highly limited by the spatial constraints imposed by the manipulators. Furthermore, the success rate of a whole-cell recording depends on mechanical interference, deterioration of recording quality during a prolonged experimental time and the tissue quality. These factors are aggravated when the number of pipettes is increased. Cleaning pipettes for immediate reuse enables recording of additional neurons in one experimental session without manual replacement. This technique has already been implemented for automated patch-clamp of single neurons in vivo (Kodandaramaiah et al., 2018; Kolb et al., 2016). Applying pipette cleaning to in-vitro multipatch setups has the potential to increase the number of pipettes with successful recordings and therefore the experimental yield. Sequential recordings from multiple cells using the same pipette would also overcome the limitation on maximum cluster size given by the number of manipulators in use. This will provide a more complete view of the microcircuit enabling the analysis of more complex network motifs and higher degree distributions (Perin et al., 2011; Song et al., 2005; Vegué et al., 2017).

### In this report, we introduce an in-vitro multipatch approach with an automated pipette pressure and cleaning system strongly facilitating recordings of up to 10 neurons simultaneously

We show that this approach and further optimizations increase the rate of complete clusters, decrease experimental time and enable sequential patching of additional neurons. We demonstrate that probing of up to 600 synaptic connections in human brain slices in one day with two setups is possible allowing comparisons of microcircuits between individual patients.

## Methods

We will first provide technical instructions for establishing a multipatch setup and how to address specific challenges when using up to 10 manipulators. We then describe an automated pressure device for multiple pipettes and how to implement a pipette cleaning protocol. Finally, we show that these technical improvements enable a higher data yield allowing extensive connectivity analysis in slices of human cortical resections and higher degree analysis of individual cells far exceeding the capacity of conventional multipatch setups.

### Spatial arrangement of hardware

#### Expanding a setup from a single or dual patch-clamp configuration to one with more patch pipettes is primarily an issue of spatial arrangement of the micromanipulators

This upscaling of multipatch setups has been accomplished with manipulator systems from different commercial manufacturers such as Luigs & Neumann (Guzman et al., 2016; Jiang et al., 2015; Wang et al., 2015; Winterer et al., 2017) and Scientifica (Cossell et al., 2015; Peng et al., 2017). Crucial to any multipatch setup is the fine spatial arrangement of the manipulators which we found most easy to address by using a custom stage even though there are also commercial solutions available. We arranged ten Sensapex uMp-Micromanipulators on a setup with a Nikon Eclipse FN1 microscope using a custom-made stage which was fabricated by a common metal workshop (Fig. 1A-C). We also successfully established a setup with 8 Scientifica PatchStar manipulators on a custom-made stage with a Scientifica SliceScope microscope (Fig. 1D, (Peng et al., 2017)). Our custom stages consist of a 10 mm aluminium sheet for stability glued to a 1 mm stainless steel sheet to create a ferromagnetic surface for cable routing using magnets. The stage is supported by platforms mounted onto poles, which enable flexible height adjustment. For exact dimensions of the stage, see supplementary material. The remaining perfusion system and hardware including electronic components resemble other typical patch-clamp setups and are listed in table 1. To operate 10 patch-clamp amplifiers within one digitizer that is also commercially available (Power1401-3A from Cambridge Electronic Design), we needed to increase the number of digital-to-analogue converter (DAC) channels from 8 to 10. We achieved this by a simple routing device that is capable of switching between channels enabling four headstages to receive stimulation from only two DAC outputs, see supplementary material for details of the routing device.

**Table 1:** Hardware components of 10-manipulator setup

| Hardware | Manufacturer | Model |
| --- | --- | --- |
| 5 dual patch-clamp amplifiers | Molecular Devices | MultiClamp 700B<br>CV-7B headstage |
| Data acquisition system | Cambridge Electronic Design | Power1401-3A<br>+ Signal Expansion (2701-5) |
| 10 Micromanipulators | Sensapex | u-Mp Micromanipulator |
| Microscope | Nikon | Eclipse FN1 |
| Motorized stage for microscope | Scientifica | UMS-2550P for x-/y-axis<br>Stepper motor for z-axis drive |
| Camera | Hamamatsu | Orca-Flash 4.0 |
| LED infrared light source | Thorlabs | M780L3 |
| 4 height adjustable poles | Thorlabs | P250/M, PB1, C1515/M |
| Active vibration isolation table | Accurion | Halcyonics_i4large |
| Peristaltic pump | Gilson | Minipuls 3 |
| Thermostat | Multichannel systems | TC01 |
| Stage for manipulators | Custom-made | Design in supplement |
| Recording chamber | Custom-made | Design in supplement |

**Figure 1:**
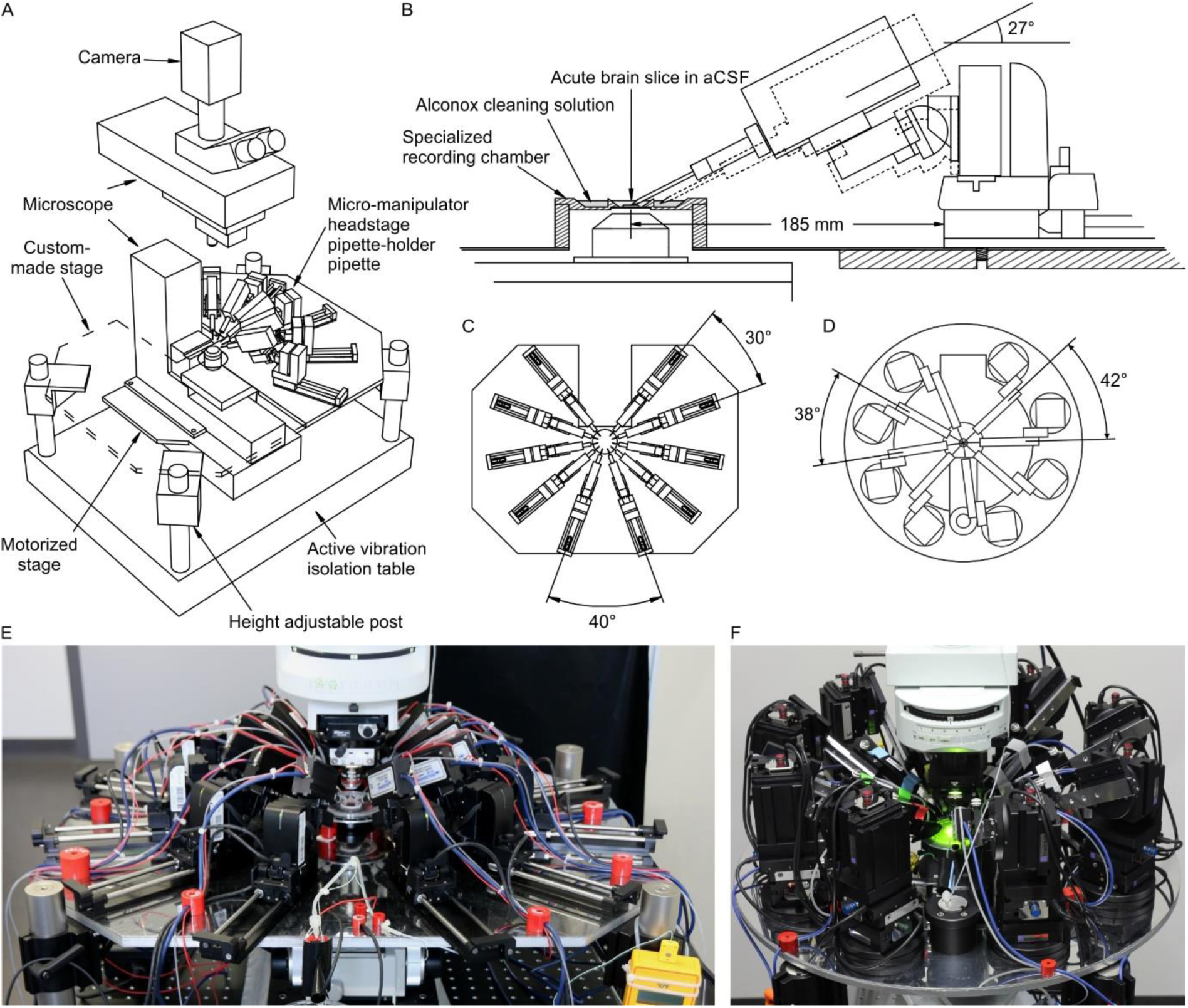
Multipatch setups. (A) Overview of essential components of a 10 -manipulator setup with Sensapex micromanipulators. (B) Side view depicting the spatial arrangement of a manipulator including headstage and pipette relative to the recording chamber which is elevated above the condensor. The dashed outline shows the manipulator with pipette t ip immersed in the cleaning solution. (C) Top view of 10 Sensapex manipulators with headstages and their angles to each other on the custom-made stage. (D) Top view of 8 Scientifica PatchStar manipulators with headstages and their angles to each other on a custom -made ring-shaped stage. (E) Photograph of the 10 -manipulator setup. (F) Photograph of the 8 -manipulator setup.

#### Another crucial aspect is the distance and angle of the pipettes to the recording chamber

(Fig. 1B). A shallow approach angle increases the range of motion in the x-axis which is important to reach both the centre of the recording well and the outside well containing the cleaning solution. However, the angle needs to be sufficiently steep to allow the pipette tip to reach the slice surface in the z-axis while avoiding touching the wall of the centre well. We found that 27° to be optimal for our custom-made recording chamber, which is positioned on an elevated platform. To increase stability, the distance of the manipulator to the recording chamber should be minimized. At the same time, sufficient space between the headstages must be maintained to allow the necessary range of motion.

### Pressure control system

#### We adapted the use of solenoid valves and pressure regulators for automated in-vivo patch-clamp to our in-vitro multipatch approach

(Kodandaramaiah et al., 2016). Throughout the process of patching a cell, variable pressure levels are required. Conventionally, this is achieved with pressurized chambers, a three-way valve and a mouthpiece or a syringe. Since control of individual pipettes using manual pressure is not feasible for multiple pipettes, an automated pressure system is needed. A previous report has demonstrated an electrically controlled pressure system for a multipatch setup (Perin and Markram, 2013). Kodandaramaiah et al. have furthermore introduced the Autopatcher, an automatic pressure device combined with a pipette movement algorithm, for automated in-vivo patch-clamp (Kodandaramaiah et al., 2016, 2012). This latter device has also been developed for multi-neuron in-vivo recordings with 4 pipettes (Kodandaramaiah et al., 2018). We adapted the described automatic pressure system to our setup and scaled it up to 10 pipettes, with different components that are cost-efficient and widely available. It also enables the application of multiple pressure levels to individual pipettes simultaneously. A detailed component list with price estimates and construction manual with illustrated step-by-step instructions for the automated pressure system is attached in the supplementary material. It can be built within a week with basic electrical equipment and does not require extensive technical skills. We believe that the easy-to-build modular construction approach is useful for labs that want to implement this tool and adjust it to their needs.

#### Multiple pressure regulators were connected to the compressed air outlet to generate different adjustable pressure levels to the pipettes

(Fig. 2A). We set one pressure regulator at 20 mbar (LOW) for continuous outflow of pipette solution during the idle state in the bath to prevent clogging of the tip. We apply 70 mbar (HIGH) for moving through the slice to approach the cell. Since HIGH pressure is only applied to the active pipette and will always be followed by the pressure of a mouth piece or syringe (PATCH), these two pressure levels share one output. In the PATCH mode we wait for the formation of the gigaseal, apply light suction when needed and apply a stronger suction to break into the cell. We believe that this is a crucial part of the patching process which needs fine manual adjustment regarding the duration and strength of suction to achieve maximum success rate (up to 88% on 6-manipulator setup, see results on performance below) while reports on fully automated in-vitro patch-clamp algorithms have reported a relatively low success rate of 43% using a single pipette (Wu and Chubykin, 2017). After successful establishment of a whole-cell configuration, the pressure in the pipette will be switched to atmospheric pressure (ATMOSPHERE). The fifth pressure channel is implemented for the pipette cleaning process described below (CLEAN).

**Figure 2:**
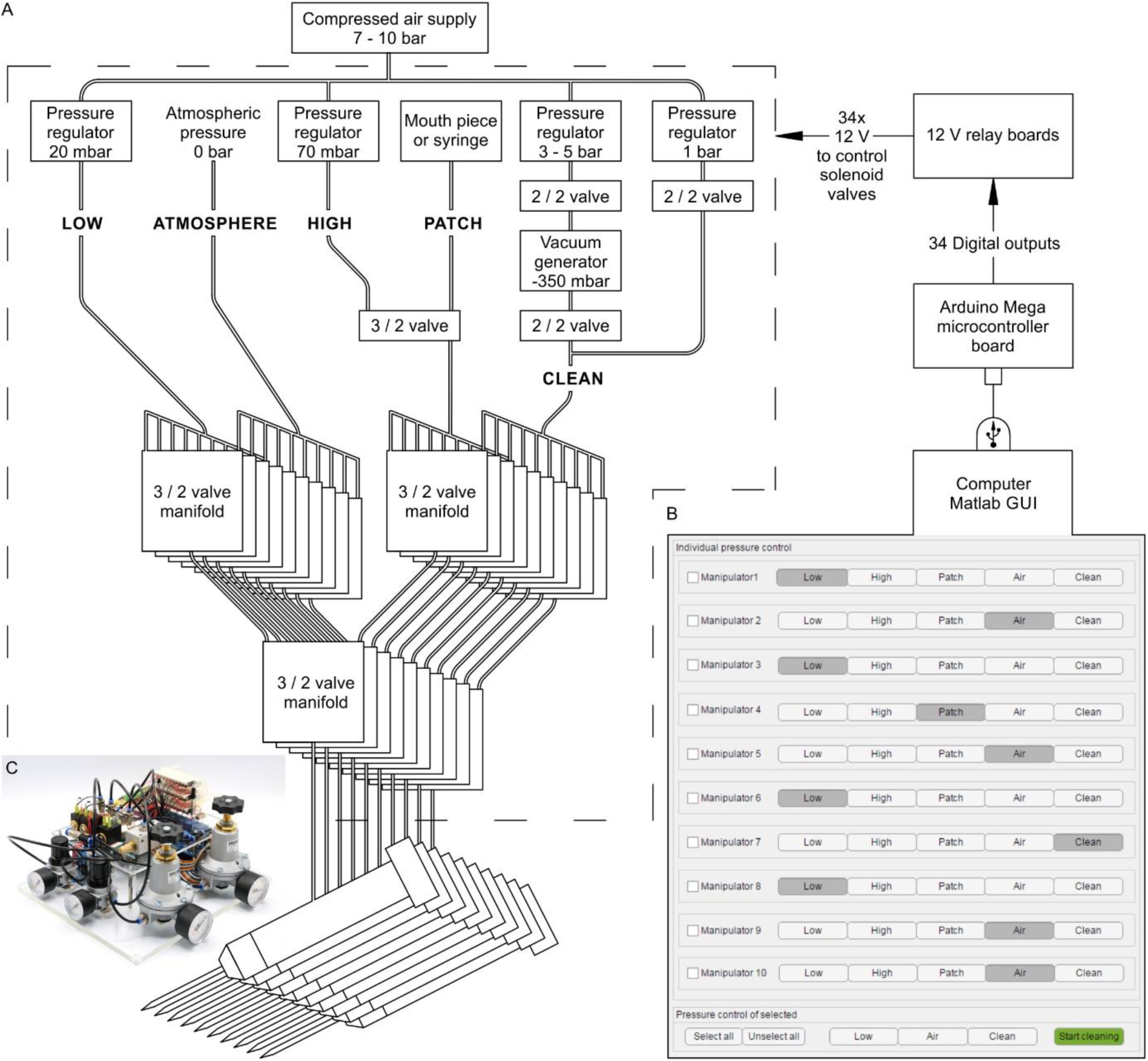
Automated pressure system. (A) Tubing scheme between the different pneumatic components. Top row depicts pressure regulators and their set pressure for each path. 2/2 valves are solenoid valves with two ports and two positions (open or closed). 3/2 valves are min iature solenoid valves with three ports and can switch between two positions connecting either inlet to one outlet. For further details, see the supplementary document. (B) Screenshot of Matlab GUI controlling the pressure system. It communicates with an A rduino microcontroller board which in turn has digital outputs connecting onto a relay board. For further details on GUI and wiring scheme see the supplementary document. (C) Photograph of an assembled pressure system.

By arranging 3 miniature solenoid valves in a tree structure for each pipette, we are able to direct these different pressure levels to each pipette individually (Fig. 2A). The solenoid valves are electrically operated using commercially available 12 V relays which are controlled by the digital outputs of an Arduino Mega microcontroller board. To control the pressure system through the Arduino board we developed a graphical user interface in MATLAB (Fig 2B, further instructions and link to code repository provided in supplementary material). This open-source and affordable pressure system is applicable to any other patch-clamp setup. It can reduce experimental errors, increase the experimental speed and help the experimenter to concentrate on the other tasks which are essential for successful operation of a multipatch setup.

### Cleaning system

#### We adopted a pipette cleaning protocol to the in-vitro multipatch setup which enables immediate retry after a failed patch attempt and recording of additional neurons through sequential patching

A recent study has shown that dipping the tip of a patch pipette in a detergent solution (Alconox) and applying a sequence of pressure and suction could clean a pipette for immediate reuse (Kolb et al., 2016). In this study, they showed that more than ten successful “repatches” using the same patch pipette were possible with no change in the electrophysiological properties of the cells were observed. We implemented this protocol to our multipatch approach, because it could increase the success rate of establishing good recordings. To incorporate a well for the detergent solution, we constructed a custom recording chamber with an outer circular well which is separated to the inner recording well (Fig 3A, see supplementary material for detailed construction designs). With the optimal angle and positioning of the manipulators (Fig. 1B), all pipettes can access the brain slice and the cleaning solution without mechanical interference.

**Figure 3:**
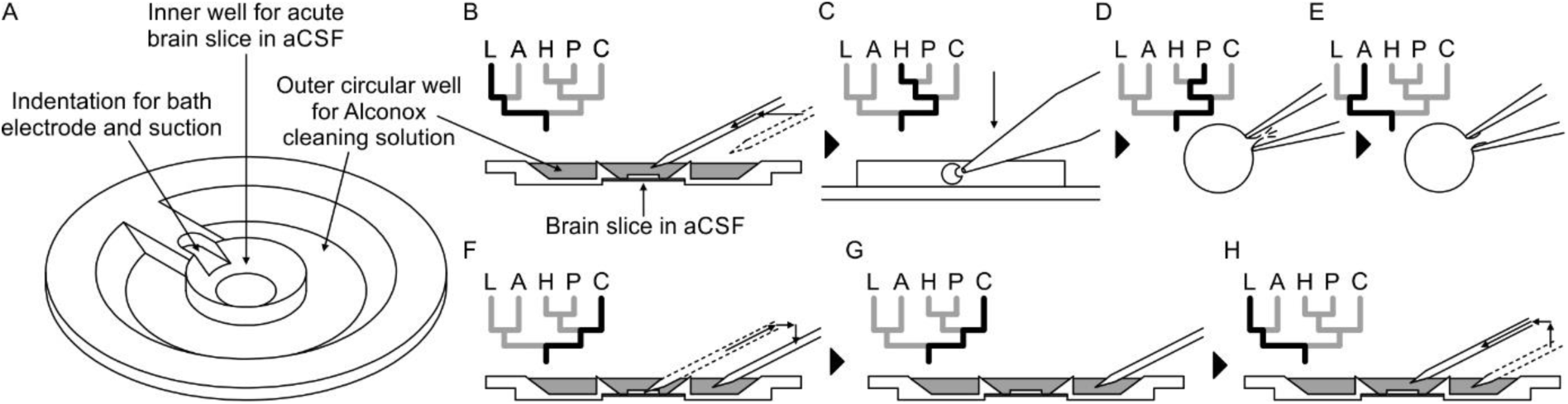
Pipette cleaning protocol. (A) Design of custom-made recording chamber with one centre well for the brain slice and recording solution and another circular outer well for the cleaning solution, see construction design in supplementary document. (B -H) Experimental steps for pipette cleaning. Tree diagrams on the top left depict the configuration of the pressure system with the following pressure levels: LOW, ATMOSPHERE, HIGH, PATCH, CLEAN. (B) Pipettes are moved into the recording solution with LOW pressure. (C) Cells are approached w ith HIGH pressure. (D) Formation of gigaseal and membrane rupture with pressures applied through the PATCH channel with either mouth piece or syringe. (E) Cells are kept at ATMOSPHERE pressure after successful patch. (F) Pipettes are moved into the cleanin g solution and switched to CLEAN pressure. (G) Cleaning pressure sequence is applied in the outer well. (H) Pipettes are moved back into the recording solution above the slice and switched to LOW pressure.

#### To clean the pipettes automatically, both the sequence of pipette pressure and manipulator movement need to be automated und coordinated

We programmed the movements of our Sensapex and Scientifica micromanipulators in Matlab and integrated them into the GUI of the pressure system. We recommend custom development of these manipulator movements due to the specific parameters of each setup. Example code and key aspects of manipulator programming can be found in the Github repository (link provided in supplementary material). Using the CLEAN channel, the automated pressure system is now able to apply positive and negative pressure as described in the protocol (Kolb et al., 2016). We used a pressure regulator connected to the compressed air outlet to generate 1 bar for expulsion and another pressure regulator coupled with a vacuum ejector to generate -350 mbar for suction at the pipette tip (Fig 2A). The pressure regulators and the vacuum generator are connected to downstream solenoid valves which control and alternate the pressure.

#### Incorporating automated manipulator movements, pressure control and the cleaning protocol make the experimental steps easier and faster (Video 1)

Pipettes are sequentially moved into the recording well to their dedicated position in the field of view approximately 2 mm above the slice. Small manual adjustments under visual control are necessary for each pipette due to variation in pipette length. The pipette pressure is set to LOW (20 mbar) to avoid clogging of the tip (Fig. 3B). To speed up this process and avoid collisions, we semi-automated it by developing a pipette finding algorithm which we discuss further below. After positioning of all pipettes around approximately 200 µm above the slice, cells are approached and patched sequentially under visual guidance. To prevent contamination of the pipette tip while moving through the slice, the HIGH pressure (70 mbar) is applied to the pipette (Fig. 3C). When a dimple on the cell surface can be seen, the pressure is released by switching to the PATCH channel. Manual application of suction either through a mouth piece or manually through a syringe is then needed to obtain a good seal and to break through the membrane to establish the whole-cell configuration (Fig. 3D). Pipettes with successfully patched cells are switched to atmospheric pressure (ATMOSPHERE, Fig. 3E). If the sealing process is unsuccessful or the whole-cell recording deteriorates, even multiple pipettes can now be cleaned simultaneously through an automated process. The cleaning process starts with the automated retraction of the pipettes to a position outside of the chamber above the outer well (Fig. 3F). They are then lowered into the detergent solution (Fig. 3G). After the pipette pressure is switched to the CLEAN channel, suction (−350 mbar, 1 s) and pressure (1 bar, 1 s) are alternatively applied for 5 cycles, followed by a long expulsion sequence (1 bar, 10 s). For the next step, the pipettes are moved to the outer rim of the recording well into the recording solution for a final expulsion sequence (1 bar, 10 s) and are then moved to the initial position above the slice while the pressure is set to LOW (Fig. 3H).

#### The final expulsion sequence does not require additional wells containing aCSF as compared to the original protocol (Kolb et al., 2016)

By omitting a second well containing aCSF we ensured access of all pipettes to the cleaning solution. However, the final expelling sequence could introduce traces of Alconox solution into the extracellular recording solution. Alconox contains 10-20% sodium linear alkylbenzene sulfonate (LAS), which affects Glycin-, GABAA- and GluR6-mediated currents at concentrations around 0.001% (Machu et al., 1998). Since it has already been shown that no significant amount of LAS could be detected in the pipette solution after the cleaning cycles (Kolb et al., 2016), the remaining potential source of contamination is residual detergent adhering to the outside surface of the pipette tip. A volume of at least 25 µl Alconox would be needed to reach the critical LAS concentration of 0.001% in the extracellular solution, assuming a recording bath volume of 1 ml and a LAS concentration of 20% in the 2% Alconox solution. We estimate the adherent Alconox solution on a pipette tip to be less than 0.2 µl considering that the total volume of intracellular solution inside a pipette is usually less than 15 µl. We thus believe that the potential impact on the synaptic connectivity and electrophysiological properties is negligible considering this high degree of dilution and the continuous flow of extracellular solution at more than 2 ml/min. Likewise, we found no difference in recording quality or electrophysiological properties between pipettes at first use and after cleaning.

The entire cleaning process takes approximately 1 minute and can be executed for multiple pipettes at the same time and simultaneously to a patch attempt. Therefore, pipettes failing to establish a good recording can be subject to immediate cleaning and reuse. This greatly increases the success rate of obtaining recordings and thus the yield of a single experiment. Furthermore, after recording of a full cluster, multiple pipettes can be cleaned, and additional cells can be patched and included in the analysis. We also believe that the possibility to repetitively perform patch attempts without manual replacement of the pipettes increases the speed of learning and performance of novel experimentalists, thus lowering the barrier to establish multipatch setups.

### Preparation of pipettes

#### With increasing number of pipettes, preparations take up significant experimental time which we addressed by additional optimizations such as a multi-pipette filling device and semiautomated positioning of pipettes

Before the pipettes are positioned above the slice and are ready for patching, multiple time consuming preparatory steps are necessary, such as pipette pulling, filling and positioning. We therefore designed a device which can hold and fill multiple pipettes at the same time (see supplementary material for construction designs). It uses the prefill approach to suck intracellular solution through the tip. Backfilling using a syringe might still be necessary when prefilling is not sufficient. This approach reduced the time to fill and mount all pipettes to 11-13 minutes, while the time of during prefilling can be used for other preparations (Fig 4C). We also optimized the procedure to find and position the pipettes in a new region of interest (Video 2). Manipulator coordinates were matched to the vmicroscope coordinates using a rotation matrix and anchored to a common reference point (Perin and Markram, 2013). A semi-automated algorithm determines optimal positions of each pipette and all pipettes are moved to roughly 200 µm above the slice. The specific steps are explained in detail in the supplementary material. This semi-automated approach reduces the risk of breaking pipette tips and the time needed to complete pipette positioning to 7-9 minutes (Fig. 4C).

**Figure 4:**
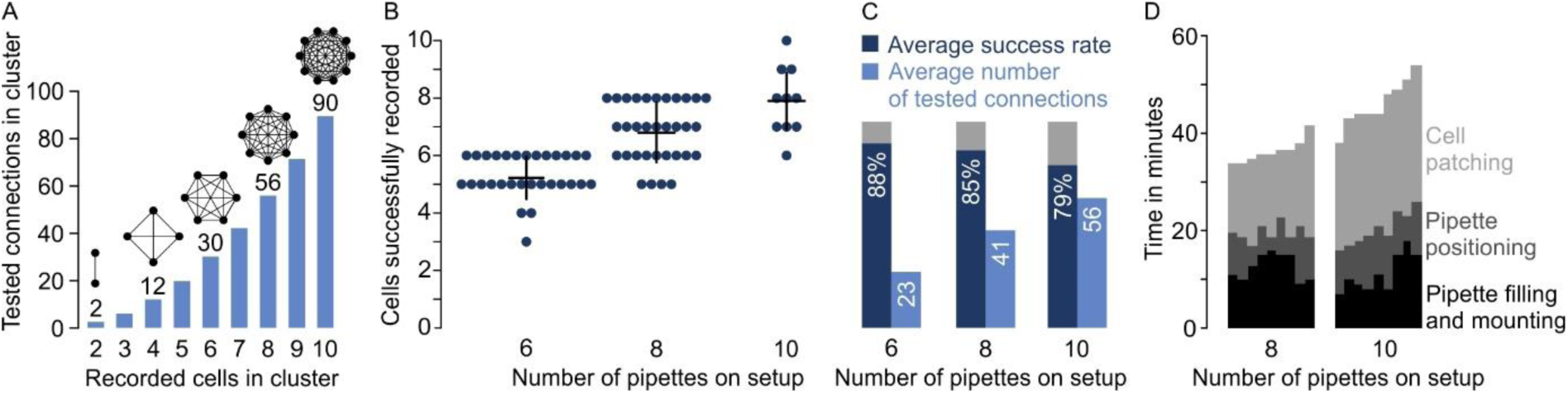
Performances on multipatch setups. (A) Bar graphs depicting maximum number of testable connections for increasing number of simultaneously recorded cells. (B) Dot plot of number of simultaneously recorded cells from rodent brain slice experiments using different multipatch setups. Black crosses indicate mean and standard deviation. Source data in Figure 5 - source data. (C) Bar graphs of performance parameters derived from B. Average success rate represents the average size of recorded clusters in relation to the number of available pipettes on each setup. Average number of tested connections are derived from the number of tested connections from each experiment. Note the decrease in success rate and the slowing increase of tested connections when the number of pipettes are increased. (D) Stac ked bar graph of time needed for individual preparatory steps of single experiments fo r the 8-manipulator setup (n=9) and the 10-manipulator setup (n=9). Source data in Figure 4 - source data.

## Results

### Performances achievable with multipatch setups

#### The number of tested synaptic connections (c) scales with more simultaneously recorded neurons (n) according to *c* = *n* × (*n -* 1) **(Fig. 4A)**

However, increasing the number of neurons lowers the rate of obtaining a full cluster with successful recordings on all pipettes, even for skilled experimenters. This is not only due to more patch attempts, but also due to the increased risk of losing established recordings through mechanical interference or deterioration of recording quality during the prolonged total time of approaching and patching cells. To quantify this observation and provide an estimate for the performance of different setup configurations, we analysed our multipatch experiments on rodent slices from previous and ongoing studies without pipette cleaning (Böhm et al., 2015; Peng et al., 2017). For the performance analysis, we focused on rodent experiments to present an applicable use case for other labs which are not working with human slices.

On a setup with 6 manipulators we obtained on average 5.3 ± 0.7 successful recordings per cluster with an average of 23 ± 7 tested connections (mean ± standard deviation, n = 30). This corresponds to an average success rate of 88% ± 12% which is, calculated as the ratio of successfully recorded cells to the maximum number of pipettes available per cluster (Fig. 4B). We refrained from manual replacement of failed pipettes due to the risk of losing the other recordings. After scaling up to 8 pipettes, we could record from 6.8 ± 1 cells on average (success rate 85% ± 13%) which increased the mean number of tested connections to 41 ± 13 (n = 33). On our setup with 10 pipettes, we recorded on average from 7.9 ± 1.1 neurons with 56 ± 17 tested connections which represents a further drop of the success rate to 79% ± 11% (n = 10). Concurrently, the total experimental time needed before the start of recording the electrophysiological properties of the neurons rose from 36.6 ± 2.3 minutes (8 pipettes) to 46.1 ± 4.6 minutes (10 pipettes), mostly due to increasing time needed to approach and patch the cells (16.8 ± 2.7 min vs 25.8 ± 2.2 min, Fig. 4D). Considering this drop in success rate and increased experimental time, the upscaling of a conventional multipatch setup becomes impractical at a certain point.

### Pipette cleaning strategies for multipatch experiments

#### The automated pipette cleaning system can be applied in different ways to improve multipatch experiments

As demonstrated above, multipatch setups will often fail to achieve their full potential due to a decreasing success rate with increasing pipette numbers. The automated pressure and cleaning system can solve this problem and increase the yield through more complete cluster recordings. Pipettes from failed recording attempts can be subject to immediate cleaning and reuse, even simultaneously to ongoing patch attempts with the next pipette (Fig. 5A). This clean-to-complete strategy increased the average cluster size on our setups with 8 and 10 pipettes to 7.8 ± 0.4 (97% ± 5% success rate, n = 16) and 9.2 ± 0.6 (92% ± 6% success rate, n = 9), respectively (Fig. 5B). After recording of a full cluster, the cleaning system can be used to probe synaptic connections to cells outside of the initial cluster (clean-to-extend, Fig. 5D). Multiple pipettes can be selected and cleaned to establish new patch-clamp recordings with other cells of interest (Fig. 5E). This clean-to-extend approach allows screening of many additional synaptic connections with minimal time investment. The number of additionally probed connections (c_new_) is dependent on the number of newly patched (n_new_) and maintained neurons (n_old_): *c*_*new*_ = 2 × (*n*_*new*_ × *n*_*old*_) + *n*_*new*_(*n*_*new*_ *–* 1). Using both strategies during rodent experiments on the 8-pipette setup with 2 to 3 recorded slices per animal, we could increase the number of tested connections from 140 ± 24 to 244 ± 52 on average per animal (Fig. 5D, mean ± standard deviation, n = 6 animals). Clean-to-extend can also be used to selectively patch more cells of interest while a set of cells are maintained. This would enable exploration of specific connectivity patterns and a more elaborate degree analysis of specific cells.

**Figure 5:**
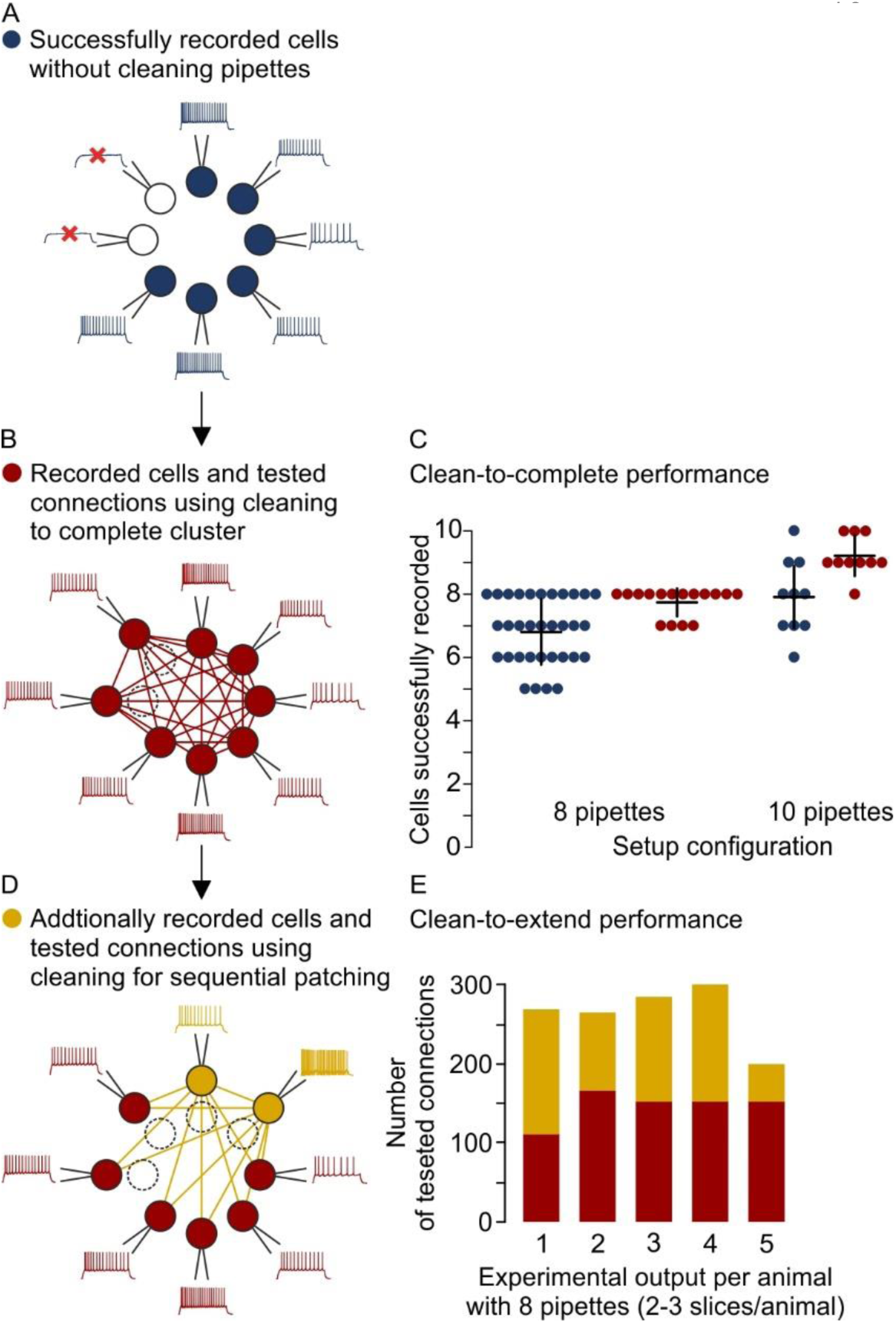
Different strategies for cleaning pipettes. (A) Scheme of successful whole-cell recordings on six pipettes (blue circles with action potentials) and failed patch attempts on two pipettes (white circles without action potentials). (B) Clean-to-complete: The two failed pipettes were cleaned and successful recordings were established from two neighbouring cells. Red circles represent a cluster that has been subject to pipette cleaning after failed patch attempts and the red lines indicate the tested synaptic connections. (C) Dot plot comparing the number of successfully recorded neurons in clusters without cleaning (blue) and with cleaning (red). Black crosses indicate mean and standard deviation. See Figure 5 -source data 1. (D) Clean-to-extend: After recording of a cluster, individual pipettes can be cleaned and used to patch additional cells (yellow circles). Yellow lines indicate the synaptic connections that could be tested due to this approach. (E) Stacked bar graph depicting the number of tested connections of the initially recorded cluster with clean-to-complete (red) and the number of additionally tested connections after clean-to-extend (yellow) from 5 animals. Data were extracted from experiments in the rat presubiculum on the 8 -manipulator setup. 2 to 3 slices were analysed from each animal in a time window of 5 to 6 h. See Figure 5 - source data 2.

#### Combining multipatch setups with the optimizations and cleaning strategies allows for extensive and efficient analysis of human microcircuits

Due to the scarce availability of human tissue from epilepsy resection surgery, a highly efficient usage of this material is desirable. With two rounds of clean-to-extend, we were able to record electrophysiological properties and the synaptic connections of up to 17 neighbouring neurons in one human slice (Fig. 6A). In this experiment, we maintained 2 neurons (1.1 & 1.2) across all further patch attempts providing information about their out- and in-connections to the other 16 neurons, respectively. This approach can unveil complex connectivity patterns between local neurons exceeding simple pairwise statistics (Fig. 6B-D). Using two setups simultaneously, we were able to record from up to 99 neurons and thereby probe 300 to 700 potential synaptic connections from individual patients within the first 24 hours after slicing. These sample sizes acquired from single patients are comparable to the pooled dataset from 22 patients in a previous report (Seeman et al., 2018). We found excitatory connectivity between pyramidal cells in human cortical layer 2/3 ranging from 12.9% to 17.7% which is similar to those reported previously (Boldog et al., 2018; Seeman et al., 2018; Szegedi et al., 2016). This substantial improvement in multipatch experiment yield has the potential to facilitate systematic analysis of complex network properties on the level of individuals in humans and other species.

**Figure 6:**
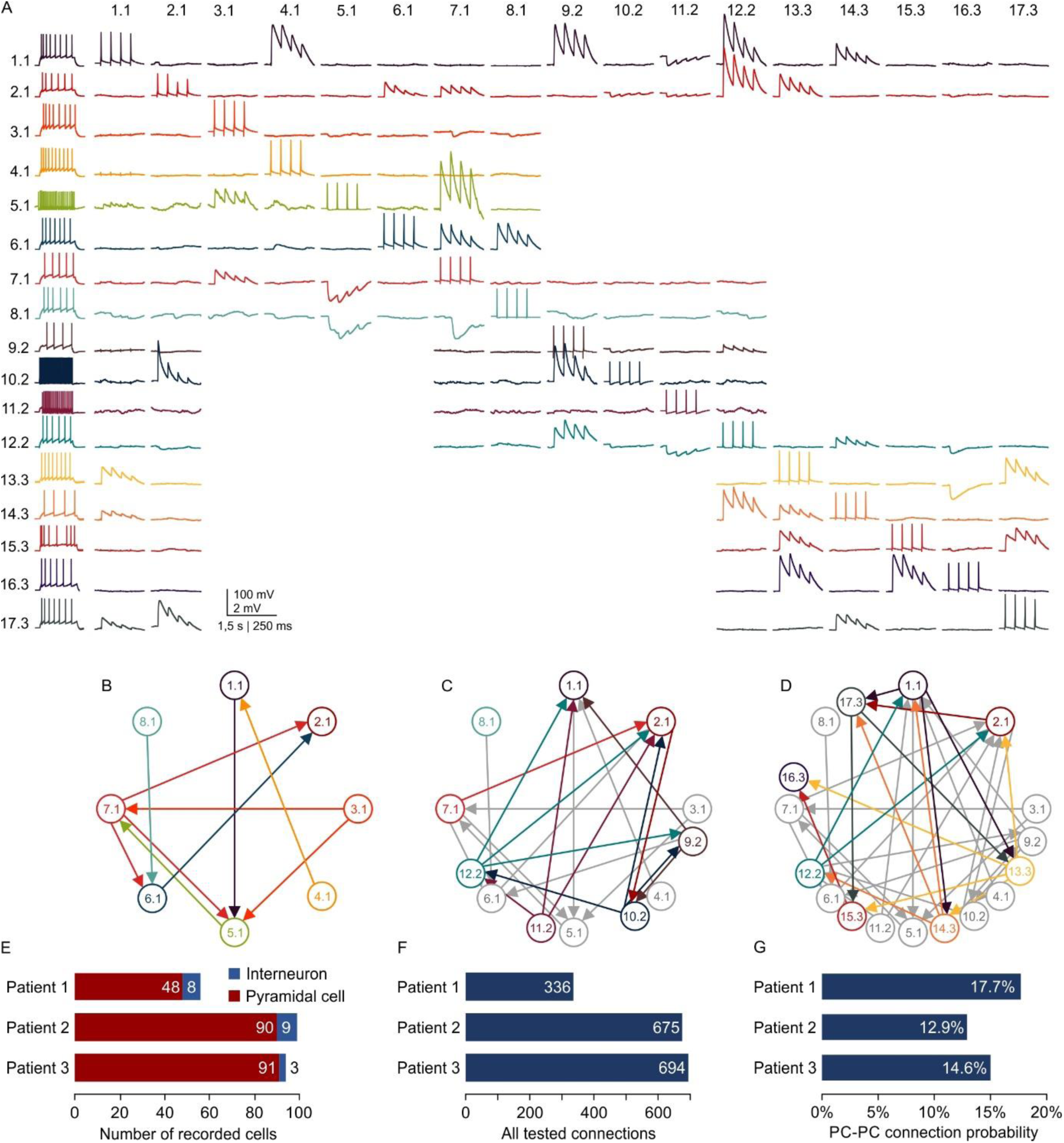
Microcircuit analysis in human slices. (A) Matrix of averaged voltage traces from 17 neurons in one acute human slice recorded on the 8-manipulator setup with two rounds of clean-to-extend. Left column shows the firing pattern of the recorded neurons. In the first session 8 neurons were patched simultaneously (cells numbered 1.1 - 8.1). Traces recorded from one cell are shown in a row with the same colour. Four action potentials were elicited in each neuron consecutively (diagonal of the matrix). The postsynaptic responses of the other neurons are aligned in the same column. After recording of the first full cluster of 8 neurons, 4 pipettes were cleaned and additional neurons in the vicinity were patched and recorded with the same stimulation protocol (9.2 - 12.2). After the second recording session, another 5 pipettes were subject to cleaning and 5 new neurons were patched, while the pipettes on neuron 1.1, 2.1 and 12.2 were not removed. This allowed screening of additional connections among the neurons from the third recording session (13.3 - 17.3), but also connections between neurons from previous recording sessions (1.1, 2.1 and 12.2). Scale bar: Horizontal 1.5 s for firing pattern, 250 ms for connection screening. Vertical 100 m V for action potentials, 2 m V for postsynaptic traces. (B) Connectivity scheme of all neurons from the first recording session with arrows indicating a detected synaptic connection. (C) Scheme of connections after first cleaning round. Coloured arrows and circles indicate neurons and connections recorded in the second session. Neurons and connections of the first recording session are shown in grey. (D) Scheme of all recorded neurons and detected connections after two cleaning rounds. Neurons and connections recorded in this third session are coloured. Neurons and conne ctions from previous recording sessions are shown in grey. In this slice, a total of 38 synaptic connections were detected out of 150 tested connections. (E) Bar graphs depicting number of recorded interneurons and pyramidal cells from three patients recorded within the first 24 hours after slicing. (F) Number of tested connections in each patient recorded within the first 24 hours. (G) Connection probability between pyramidal cells calculated by the number of found to tested connections recorded within the first 24 hours. See Figure 6 -source data.

## Discussion

### In this report, we demonstrate that equipping an in vitro multipatch setup with an automated pipette cleaning system and adding further optimizations, increases the experimental yield and generates sufficient data for microcircuit analysis within single patients

This represents a substantial increase in performance and experimental output compared to previous studies on synaptic connectivity, which usually require pooling of data across multiple subjects to achieve statistically significant results (Seeman et al., 2018; Thomson and Lamy, 2007). With 300 to 600 probed connections per patient, correlation of microcircuit properties to individual patient characteristics are now becoming feasible as has been shown for dendritic morphology and action potential kinetics (Goriounova et al., 2018). Applied to animal studies, this approach can reduce the number of animals needed to reach statistically significant findings. This is especially important when investigating animal models where ethical considerations prohibit large cohorts due to harmful interventions.

### The possibility to further explore the connectivity of neurons through clean-to-extend allows microcircuit analysis on a larger scale

While manual replacement of pipettes is possible, the risk of mechanical disruption of the recordings up to now prevented extensive practice of patching cells sequentially. Therefore, studies on synaptic connectivity have been limited by the number of available manipulators. Increasing the sample size of recorded neurons and tested synaptic connections through clean-to-extend now enables a more comprehensive analysis of the microcircuit. This is especially relevant as theoretical work has identified higher order network parameters such as triadic motifs and simplices as relevant topological constraints for microcircuit computation (Perin et al., 2011; Reimann et al., 2017; Song et al., 2005). A minimum of 3 to 4 neurons per cluster is necessary for triplet analyses, while increasing the number leads to larger sample sizes and enables higher dimensional parameters. This increase in sample size per cluster also enables a better assessment of slice to slice variability, which can help to control for methodological artefacts or confounding biological parameters. However, our impression is that increasing setups beyond 10 manipulators would come with a serious trade-off regarding experimental time and success rate. Clean-to-extend enables increasing the number of probed connections of neurons beyond this limitation which is important for network statistics such as the sample degree correlation and can be utilized for further novel graph statistics (Vegué et al., 2017).

### We addressed technical challenges in establishing and operating a highly advanced multipatch setup and provide extensive documentation on how to implement optimizations making this method easier to handle

Patch-clamp electrophysiology is a widespread and established method, but operating multiple manipulators still requires skilled researchers with long training. The low-cost and user-friendly solutions presented here can be used to upgrade existing patch-clamp setups with single or multiple manipulators. This could enable other labs to adopt the multi-neuron approach and use the optimizations to reduce experimental errors and generate results faster. Additionally, semi-automated pipette positioning as well as repetitive pipette cleaning without the need for manual replacement can help novel experimentalists to focus on the crucial steps of cell patching and may reduce the training time.

### While fully automated in-vivo patch-clamp experiments are valuable due to limited visual information for manual patching (Kodandaramaiah et al., 2018, 2012; Suk et al., 2017), the advantage of fully automated in-vitro patch-clamp is less clear

We found that manual adjustments during the visually guided approach of the cell and the fine-tuned application of pressure to obtain a whole-cell configuration yielded higher success rates (88% / 85% / 79% on 6-/8-/10-manipulator setup) compared to those reported using a fully automated system which reported a success rate of 43% using one pipette (Wu et al., 2016). Our semi-automated pipette positioning approach was faster (53 / 55 s per pipette on 8-/10- manipulator setup) than the fully automated method (103 s using one pipette) while the time needed to manually approach the cell and establish a whole-cell configuration was slightly higher (126 / 154 s per pipette on 8-/10-manipulator setup vs 120 s). Further optimized algorithms with better performance could be combined with pipette cleaning to improve the success rate and speed of multipatch experiments. However, we currently believe that fully automated multipatch experiments would require a disproportionate amount of effort for the resulting increase in performance. Also, slice exchange and cell selection would still require human presence.

### Compared to other methods, the multipatch approach provides superior access to cellular and synaptic properties with a trade-off regarding the size of the sampled network

All-optical approaches using channelrhodopsins to stimulate individual neurons and fluorescent calcium or voltage indicators to measure their responses can be used to probe connectivity of a higher number of cells than the multipatch approach (Emiliani et al., 2015). Simultaneous optical stimulation and recording using calcium indicators have been performed in up to 20 neurons (Packer et al., 2014). However, calcium transients cannot reach single action potential or subthreshold resolution, potentially missing subthreshold monosynaptic connections. Voltage indicators can resolve subthreshold signals with higher temporal resolution than calcium imaging and have been shown to detect monosynaptic connections combined with optical stimulation in organotypical slices (Hochbaum et al., 2014). But technical challenges such as imaging speed and high illumination intensities still limit its application for large network investigations in acute brain slices. While calcium imaging can also be applied in-vivo, multipatch in this condition is technically very challenging and thus far limited to a maximum of 4 pipettes (Jouhanneau et al., 2018; Kodandaramaiah et al., 2018). Patch-clamp recordings can furthermore be combined with presynaptic stimulation using extracellular electrodes, channelrhodopsin or glutamate uncaging to further map synaptic inputs from more neurons (Fino and Yuste, 2011; Jäckel et al., 2017; Wang et al., 2015). These approaches do not resolve electrophysiological and anatomical properties and further connectivity of the presynaptic neurons, but they can be used to complement the multipatch approach. Finally, multipatch experiment do not need genetically modified cells and can be applied on brain slices from a variety of species.

### Taken together, the experimental advances presented here enables highly efficient extraction of microcircuit parameters from human cortical tissue even allowing comparisons between individual patients

The potential to analyse reciprocal connections of more than 10 neurons in one cluster will help to develop more sophisticated subgraph metrics, strengthening the inference onto the underlying microcircuit. Finally, this versatile method can be applied to various species to uncover overarching principles of microcircuit topology.

## Materials and methods

### Human and animal tissue

Human tissue was acquired from temporal lobe resections of patients with pharmacoresistant temporal lobe epilepsy. All patients gave a written consent for the scientific use of the resected tissue. All procedures adhered to ethical requirements and were in accordance to the respective ethical approval (EA2/111/14).

Performance data of rodent multipatch experiments without cleaning were extracted from previous studies on mouse subiculum (6-pipette setup; Böhm et al., 2018) and rat presubiculum (8-pipette setup; Peng et al., 2017). Performance data with cleaning on the 8-manipulator setup were extracted from unpublished rat presubiculum experiments and all performance data on the 10-manipulator setup were extracted from unpublished rat motor cortex experiments. Experimental times were documented during a subset of experiments. Acute brain slices were prepared from 19 to 35 days old transgenic Wistar rats expressing Venus-YFP under the VGAT promotor (Peng et al., 2017; Uematsu et al., 2008) or from 21 to 42 days old male C57BL/6N mice (Böhm et al., 2015). Animal handling and all procedures were carried out in accordance with guidelines of local authorities (Berlin, [T0215/11], [T0109/10]), the German Animal Welfare Act and the European Council Directive 86/609/EEC.

### Slice preparation

Experimental procedure on human tissue was as previously described (Lehnhoff et al., 2019). In brief, tissue samples were collected at the operating theatre and transferred to the laboratory within 30 to 40 minutes in cooled sucrose based aCSF enriched with carbogen (95% O2, 5% CO_2_). They were cut in ice-cold sucrose aCSF containing (in mM): 87 NaCl, 2.5 KCl, 3 MgCl_2_, 0.5 CaCl_2_, 10 glucose, 75 sucrose, 1.25 NaH_2_PO_4_, and 25 NaHCO_3_ (310 mOsm), enriched with carbogen (95% O2, 5% CO_2_). After removal of residual pia, the tissue was cut into 400 µm thick slices and subsequently stored in sucrose aCSF solution heated to 34°C for 30 minutes recovery. Slice thickness was 400 µm. In a subset of experiments, an antibiotic was added to the incubation solution (minocycline 2 nM). Subsequently and until recording, the slices were stored at room temperature. Whole-cell recordings were performed at 34 °C under submerged conditions, the bath chamber was perfused with an aCSF solution containing (in mM): 125 NaCl, 2.5 KCl, 1 MgCl_2_, 2 CaCl_2_, 10 glucose, 1.25 NaH_2_PO_4_, and 25 NaHCO_3_ (300 mOsm). Patch pipettes were pulled from borosilicate glass capillaries (2 mm outer / 1 mm inner diameter; Hilgenberg) on a horizontal puller (P-97, Sutter Instrument Company) and filled with intracellular solution containing (in mM): 130 K-gluconate, 2 MgCl_2_, 0.2 EGTA, 10 Na_2_-phosphocreatine, 2 Na_2_ATP, 0.5 Na_2_GTP, 10 HEPES buffer and 0.1% biocytin (290–295 mOsm, pH adjusted to 7.2 with KOH).

The rat experiments were performed as previously reported (Peng et al., 2017). After isoflurane anaesthesia and decapitation, the head was submerged in an ice-cold sucrose artificial cerebrospinal fluid solution containing (in mM): 80 NaCl, 2.5 KCl, 3 MgCl_2_, 0.5 CaCl_2_, 25 glucose, 85 sucrose, 1.25 NaH_2_PO_4_ and 25 NaHCO_3_ (320–330 mOsm), enriched with carbogen (95% O2, 5% CO2). Brain slices of 300 µm thickness were cut on a Leica VT1200 vibratome (Leica Biosystems) and subsequently stored in the sucrose aCSF solution. After a recovery period of 30 minutes at 30°C, the slices were stored at room temperature. Whole-cell recordings were performed under submerged conditions, the bath chamber was perfused with an aCSF solution containing (in mM): 125 NaCl, 2.5 KCl, 1 MgCl_2_, 2 CaCl_2_, 25 glucose, 1.25 NaH_2_PO_4_, and 25 NaHCO_3_ (310–320mOsm). The solution was enriched with carbogen (95% O2, 5% CO2) and heated to 34°C. Patch pipettes were pulled in the same way as for the human slice experiments and filled with intracellular solution containing (in mM): 130 K-gluconate, 6 KCl, 2 MgCl_2_, 0.2 EGTA, 5 Na_2_-phosphocreatine, 2 Na_2_ATP, 0.5 Na_2_GTP, 10 HEPES buffer and 0.1% biocytin (290–300 mOsm, pH adjusted to 7.2 with KOH).

Hardware components used for visualization and electrophysiological recordings are listed in table 1. Data were low-pass filtered at 6 kHz using the built in four-pole Bessel filter and digitized at a sampling rate of 20 kHz. For synaptic connectivity screening, all cells were recorded in current clamp mode and held near –60 mV by means of constant and adjusting current injection. Four action potentials were elicited in each cell at 20 Hz with 1-4 nA for 1-3 ms and postsynaptic responses in the other cells were detected in the averaged traces from 30 to 50 sweeps. Successful recordings were defined by a sufficiently high seal-resistance, resting membrane potentials of the cells more negative than -50 mV (not corrected for liquid junction potential), long-term recording stability, and the ability of the neurons to produce characteristic action potential patterns during sustained step depolarisations.

## Acknowledgement

We are grateful to the patients for providing the tissue and thank P. Fidzinski and M. Holtkamp for clinical organization and assistance. We also thank L. Faraj for support during the creation of the pressure system assembly guide and I. Vida for providing the transgenic rats. We thank the mechanical workshop of the Charité-Universitätsmedizin Berlin for technical assistance and fabrication of the custom-made components. We thank Jochen Winterer and Rosanna Sammons for helpful comments on an earlier version of the manuscript.

## Rich media files

### Video 1: Automated cleaning of all pipettes on a 10-multipatch setup

The main image depicts a front view of the 10-manipulator setup with all manipulators subject to automated movements during a pipette cleaning sequence. The insert on the bottom left corner shows a close-up view of the pipettes and the recording chamber. For better overview, the two front manipulators were moved aside. The insert on the bottom right corner shows the DIC microscope view with a 4× objective. At the end of the sequence all pipettes are out of focus, due to different final z-positions. The time stamp in the middle shows the elapsed time in seconds. Roughly 20 second of alternating pressure sequence was cut (second 13 to 32). The last 8 second sequence shows the pressure system while switching between pressure levels. The clicking noise is generated by the relais switches.

### Video 2: Pipette positioning algorithm

This video shows the pipette movements during the positioning phase under the DIC microscope. The first sequence is in a 5× time lapse with the 4× objective. Pipettes are moved in and out automatically, note the small manual adjustments in between. The second sequence is played in real time and shows all pipettes moving to their adjusted target position underneath the 40× objective.

## Supplementary File

### Supplementary materials

### Pressure and cleaning system

#### Overview of components

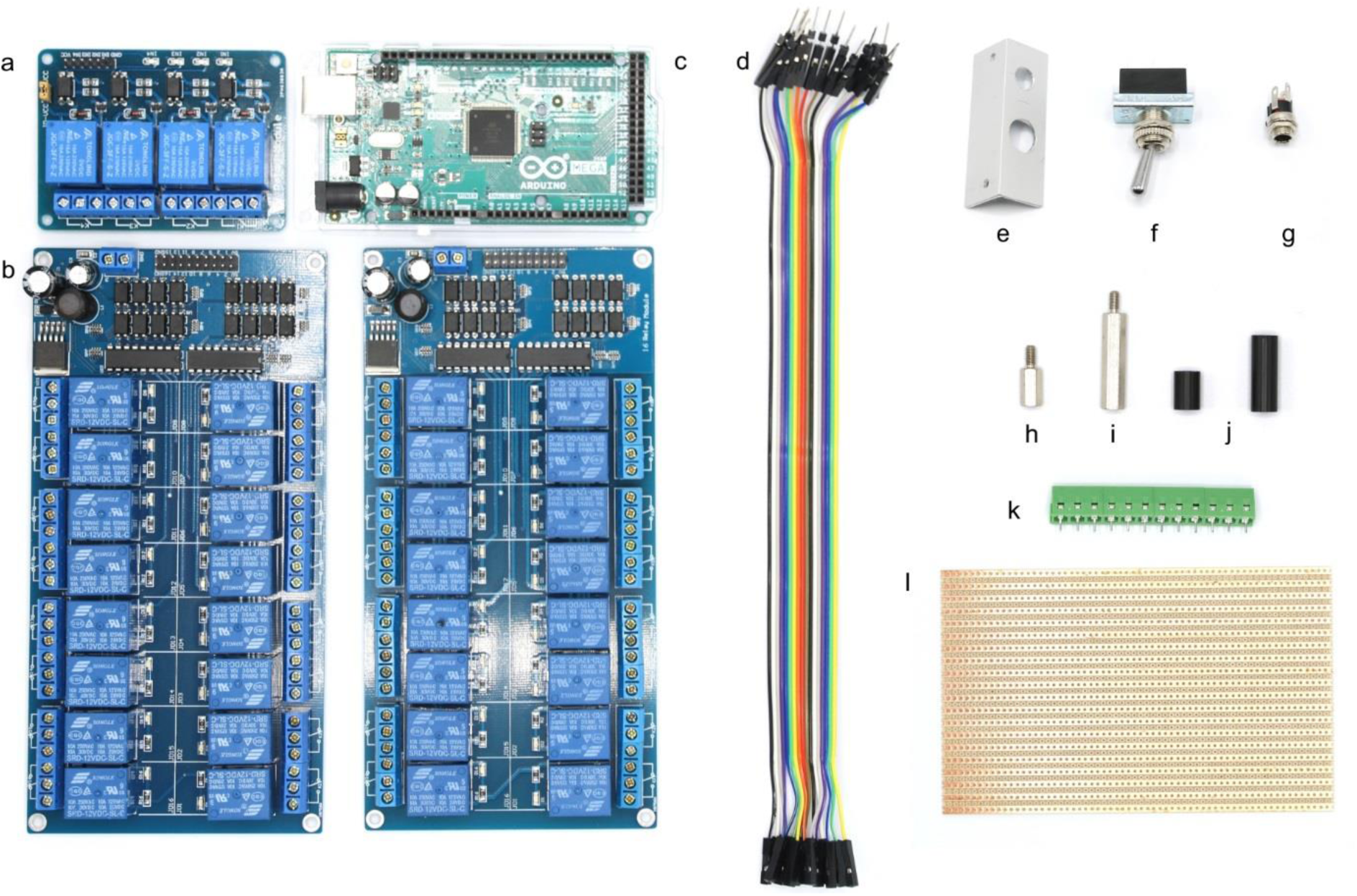

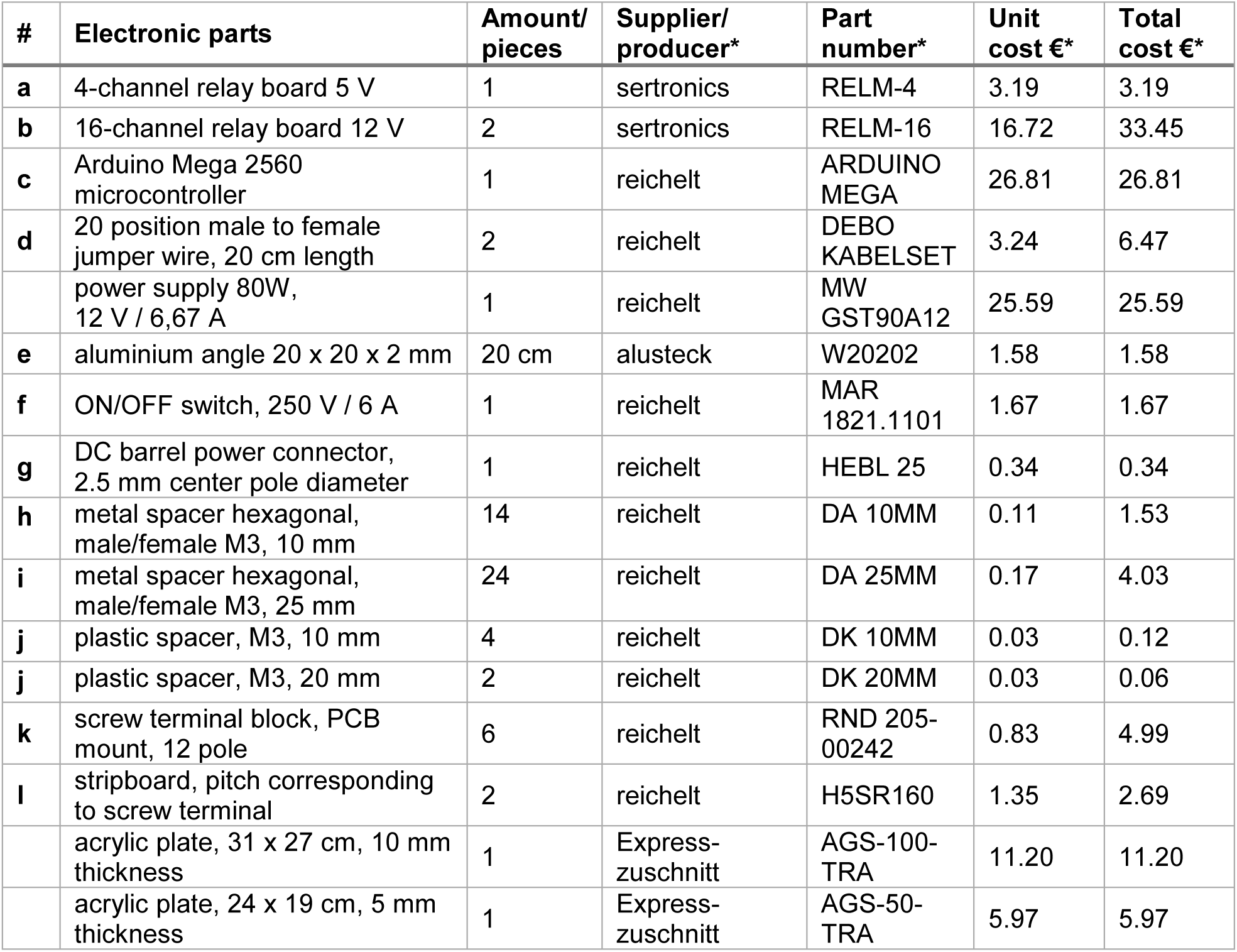
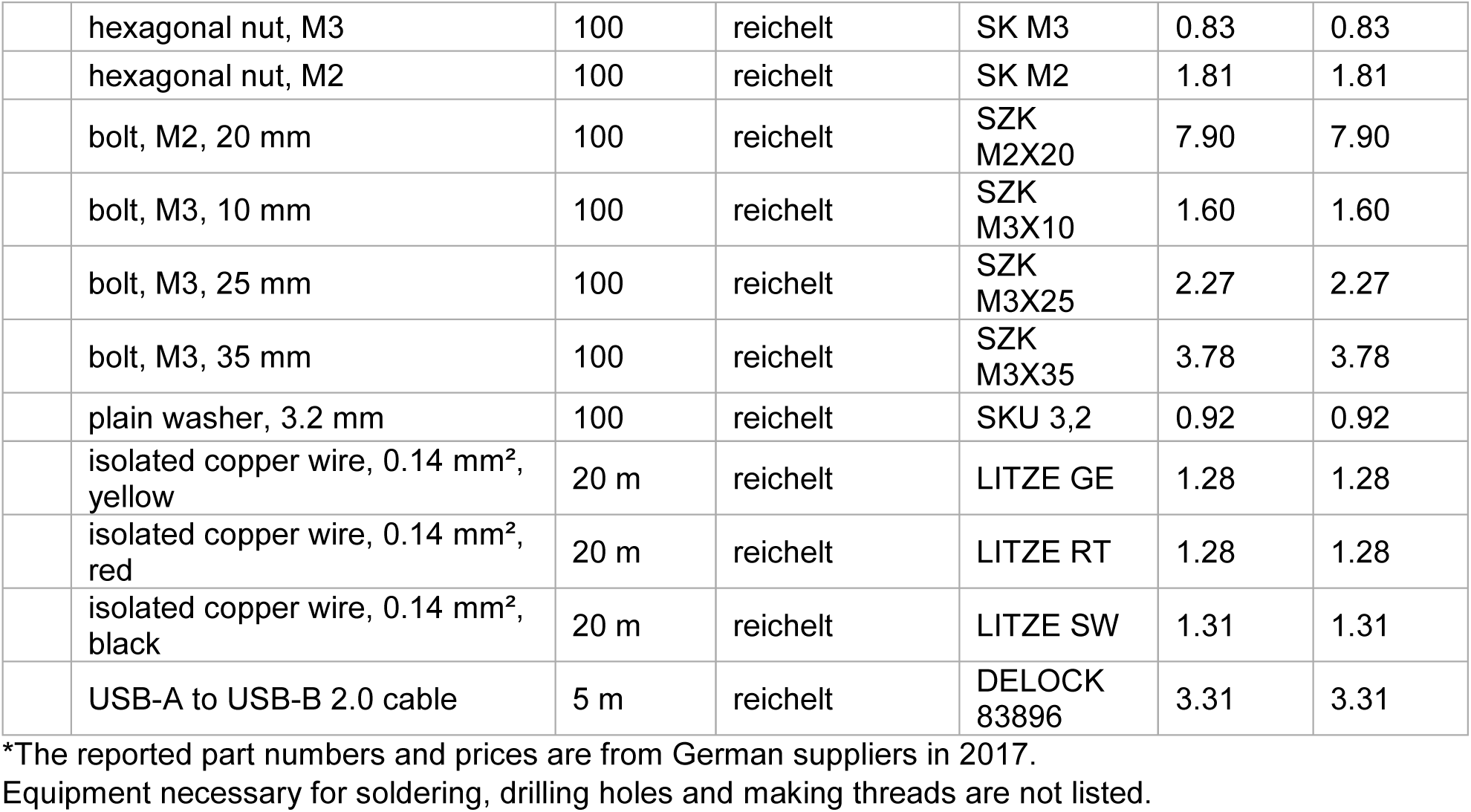

| # | Electronic parts | Amount/<br>pieces | Supplier/<br>producer* | Part<br>number* | Unit<br>cost €* | Total<br>cost €* |
| --- | --- | --- | --- | --- | --- | --- |
| a | 4-channel relay board 5 V | 1 | sertronics | RELM-4 | 3.19 | 3.19 |
| b | 16-channel relay board 12 V | 2 | sertronics | RELM-16 | 16.72 | 33.45 |
| c | Arduino Mega 2560<br>microcontroller | 1 | reichelt | ARDUINO<br>MEGA | 26.81 | 26.81 |
| d | 20 position male to female<br>jumper wire, 20 cm length | 2 | reichelt | DEBO<br>KABELSET | 3.24 | 6.47 |
|  | power supply 80W,<br>12 V / 6,67 A | 1 | reichelt | MW<br>GST90A12 | 25.59 | 25.59 |
| e | aluminium angle 20 x 20 x 2 mm | 20 cm | alusteck | W20202 | 1.58 | 1.58 |
| f | ON/OFF switch, 250 V / 6 A | 1 | reichelt | MAR<br>1821.1101 | 1.67 | 1.67 |
| g | DC barrel power connector,<br>2.5 mm center pole diameter | 1 | reichelt | HEBL 25 | 0.34 | 0.34 |
| h | metal spacer hexagonal,<br>male/female M3, 10 mm | 14 | reichelt | DA 10MM | 0.11 | 1.53 |
| i | metal spacer hexagonal,<br>male/female M3, 25 mm | 24 | reichelt | DA 25MM | 0.17 | 4.03 |
| j | plastic spacer, M3, 10 mm | 4 | reichelt | DK 10MM | 0.03 | 0.12 |
| j | plastic spacer, M3, 20 mm | 2 | reichelt | DK 20MM | 0.03 | 0.06 |
| k | screw terminal block, PCB<br>mount, 12 pole | 6 | reichelt | RND 205-00242 | 0.83 | 4.99 |
| l | stripboard, pitch corresponding<br>to screw terminal | 2 | reichelt | H5SR160 | 1.35 | 2.69 |
|  | acrylic plate, 31 x 27 cm, 10 mm<br>thickness | 1 | Express-<br>zuschnitt | AGS-100-TRA | 11.20 | 11.20 |
|  | acrylic plate, 24 x 19 cm, 5 mm<br>thickness | 1 | Express-<br>zuschnitt | AGS-50-TRA | 5.97 | 5.97 |
|  | hexagonal nut, M3 | 100 | reichelt | SK M3 | 0.83 | 0.83 |
|  | hexagonal nut, M2 | 100 | reichelt | SK M2 | 1.81 | 1.81 |
|  | bolt, M2, 20 mm | 100 | reichelt | SZK<br>M2X20 | 7.90 | 7.90 |
|  | bolt, M3, 10 mm | 100 | reichelt | SZK<br>M3X10 | 1.60 | 1.60 |
|  | bolt, M3, 25 mm | 100 | reichelt | SZK<br>M3X25 | 2.27 | 2.27 |
|  | bolt, M3, 35 mm | 100 | reichelt | SZK<br>M3X35 | 3.78 | 3.78 |
|  | plain washer, 3.2 mm | 100 | reichelt | SKU 3,2 | 0.92 | 0.92 |
|  | isolated copper wire, 0.14 mm<sup>2</sup>,<br>yellow | 20 m | reichelt | LITZE GE | 1.28 | 1.28 |
|  | isolated copper wire, 0.14 mm<sup>2</sup>,<br>red | 20 m | reichelt | LITZE RT | 1.28 | 1.28 |
|  | isolated copper wire, 0.14 mm<sup>2</sup>,<br>black | 20 m | reichelt | LITZE SW | 1.31 | 1.31 |
|  | USB-A to USB-B 2.0 cable | 5 m | reichelt | DELOCK<br>83896 | 3.31 | 3.31 |

\*The reported part numbers and prices are from German suppliers in 2017.
Equipment necessary for soldering, drilling holes and making threads are not listed.

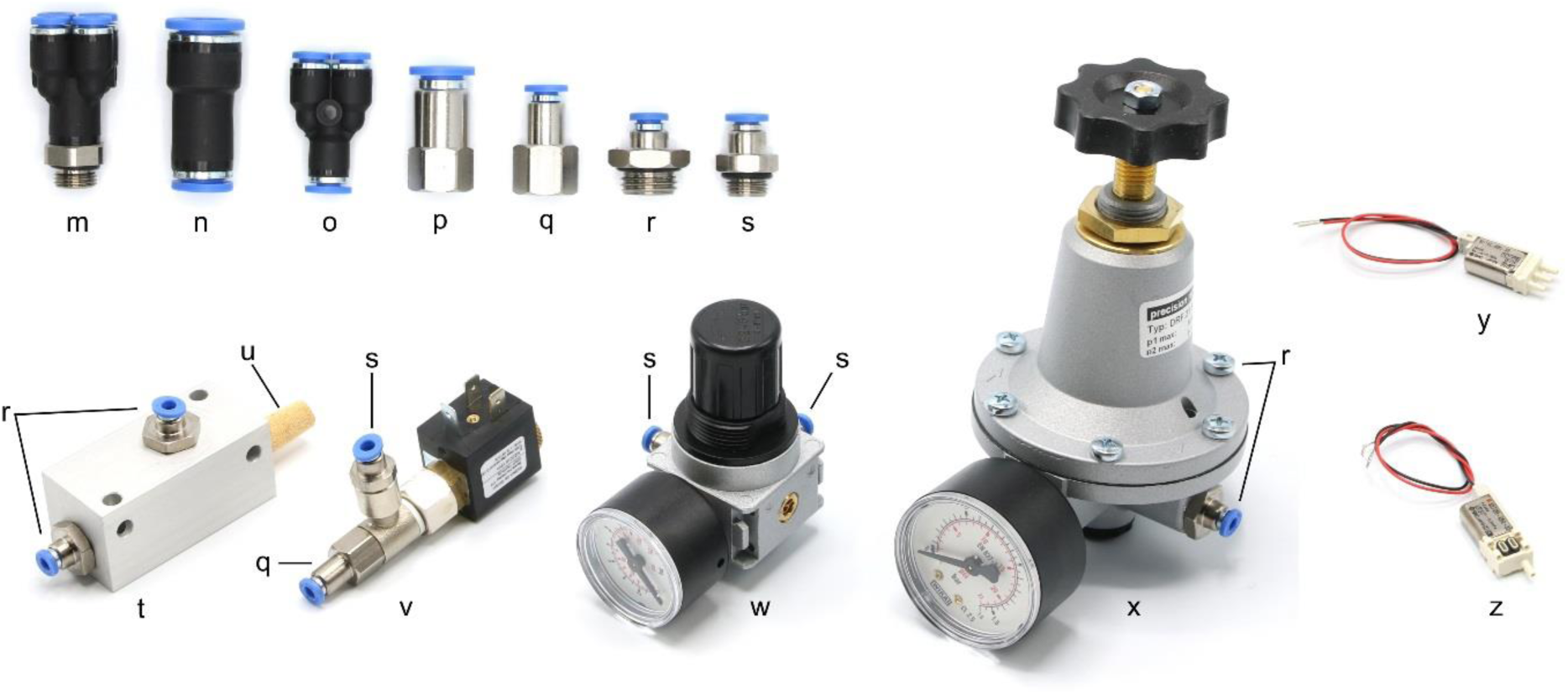

| # | Pneumatic parts | Amount | Supplier/<br>Producer* | Part number* | Unit<br>cost €* | Total<br>cost €* |
| --- | --- | --- | --- | --- | --- | --- |
| **m** | push-in fitting, multiple distributor 4 outlets, G1/8 external thread, 4mm tubing | 1 | Esska | IQSQ184G0000 | 3.29 | 3.29 |
| **n** | push-in reducing connector, 10mm/8mm tubing | 1 | Esska | IQSG10800000 | 2.25 | 2.25 |
| **o** | push-in Y-fitting, 4mm tubing | 1 | Esska | IQSY40000000 | 1.50 | 1.50 |
| **p** | push-in fitting, G1/8 internal thread, 8mm tubing | 1 | Esska | IQSF18800000 | 1.62 | 1.62 |
| **q** | push-in fitting, G1/8 internal thread, 4mm tubing | 3 | Esska | IQSF18400000 | 1.64 | 4.92 |
| **r** | push-in fitting, G1/4 external thread, 4mm tubing | 6 | Esska | IQSG144G0000 | 1.08 | 6.48 |
| **s** | push-in fitting, G1/8 external thread, 4mm tubing | 7 | Esska | IQSG184G0000 | 0.96 | 6.72 |
| **t** | vacuum ejector | 1 | Esska | 175811211 | 70.26 | 70.26 |
| **u** | silencer for vacuum ejector | 1 | Esska | SD14MS000000 | 1.06 | 1.06 |
| **v** | 2/2 solenoid valve | 3 | Esska | 8552MZ12V000 | 32.14 | 96.42 |
| **w** | pressure regulator, 1-10 bar | 2 | Esska | R018-6000000 | 29.85 | 59.70 |
| **x** | precision pressure regulator, 10-1000 mbar | 2 | Esska | DRF31GS00000 | 68.38 | 136.76 |
| **y** | 3/2 solenoid valve, body ported | 12 | SMC | S070C-6BG-32 | 38.04 | 456.48 |
| **z** | 3/2 solenoid valve, body ported manifold | 20 | SMC | S070M-6BG-32 | 34.06 | 681.20 |
|  | U end plate assembly | 2 | SMC | SS070M01-2A | 14.85 | 29.70 |
|  | d end plate assembly | 2 | SMC | SS070M01-3A | 14.85 | 29.70 |
|  | PE-tubing 50 m, outer diameter 4 mm, inner diameter 2 mm | 1 | Esska | 7031PL4X2SCH | 11.72 | 11.72 |
|  | silicone tubing 25 m, outer diameter 4 mm, inner diameter 2 mm | 1 | Roth | 9559.1 | 20.35 | 20.35 |
| **\*** | thread sealing tape | 1 | Esska | 929500064514 | 1.67 | 1.67 |

\*Use thread sealing tape to seal the connections between push-in fittings (p,q,r,s) the vacuum ejector (t), the 2/2 solenoid valves (v), the pressure regulators (w, x) and the multiple distributor (m).

#### Layout of acrylic sheet

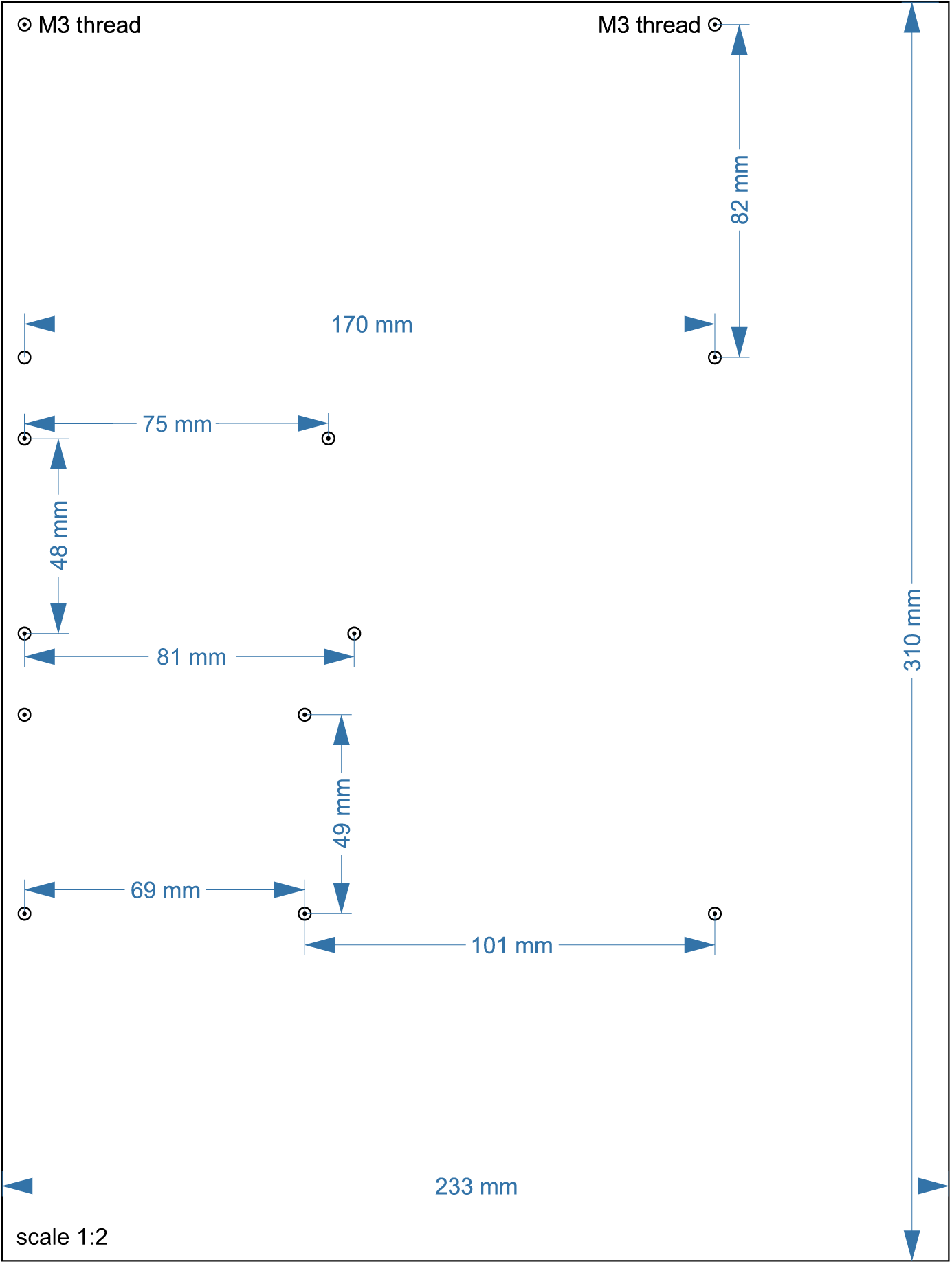

Layout of lower acrylic sheet with 10 mm thickness. Circles with central dot indicate positions where a M3 thread needs to be cut (first drill hole with a diameter of 2.5 mm). The scale is 1:2, a 1:1 enlarged layout could serve as a printed template.

#### Layout of acrylic sheet and angle

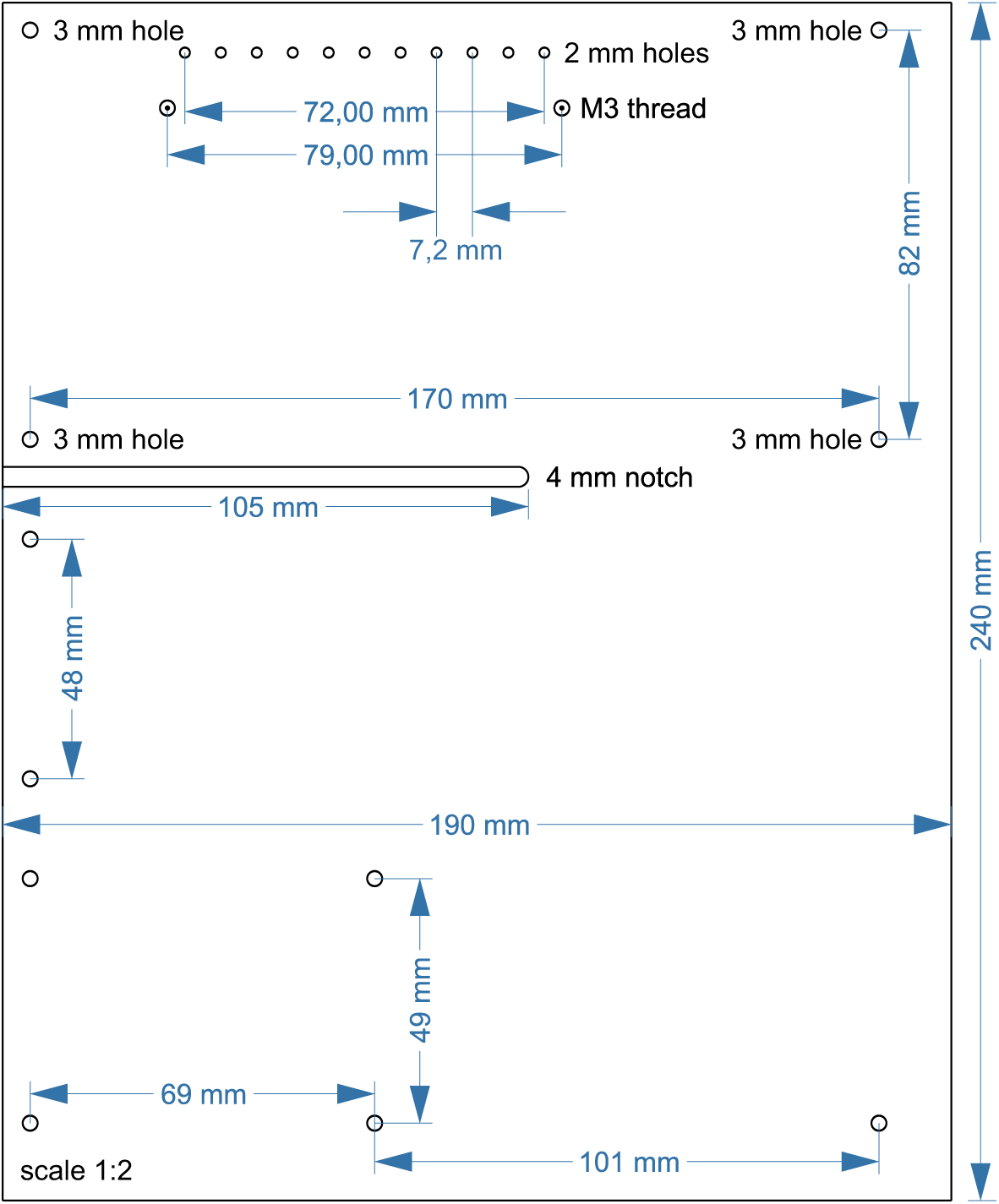

Layout of upper acrylic sheet with 5 mm thickness. Cut 2 × M3 threads at circles with central dot (first drill 2.5 mm holes). Drill 11 × 2 mm holes, each 7,2 mm apart. All other circles indicate 3 mm holes. A 4 mm notch will allow for passage of cables. The scale is 1:2, a 1:1 enlarged layout could serve as a printed template.

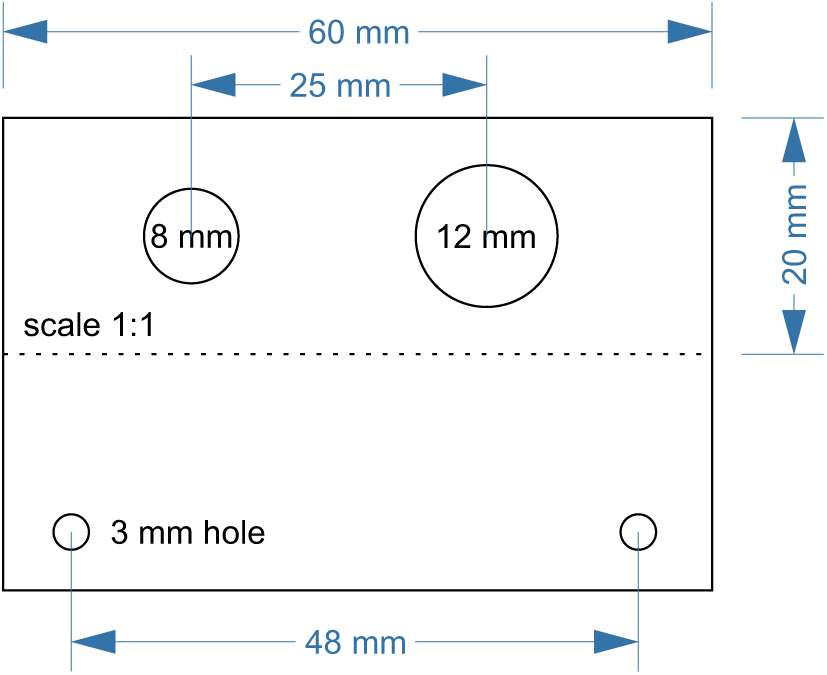

Layout of upper acrylic sheet with 5 mm thickness. Cut 2 × M3 threads at circles with central dot (first drill 2.5 mm holes). Drill 11 × 2 mm holes, each 7,2 mm apart. All other circles indicate 3 mm holes. A 4 mm notch will allow for passage of cables. The scale is 1:2, a 1:1 enlarged layout could serve as a printed template.

#### Assembly guide for pressure control device

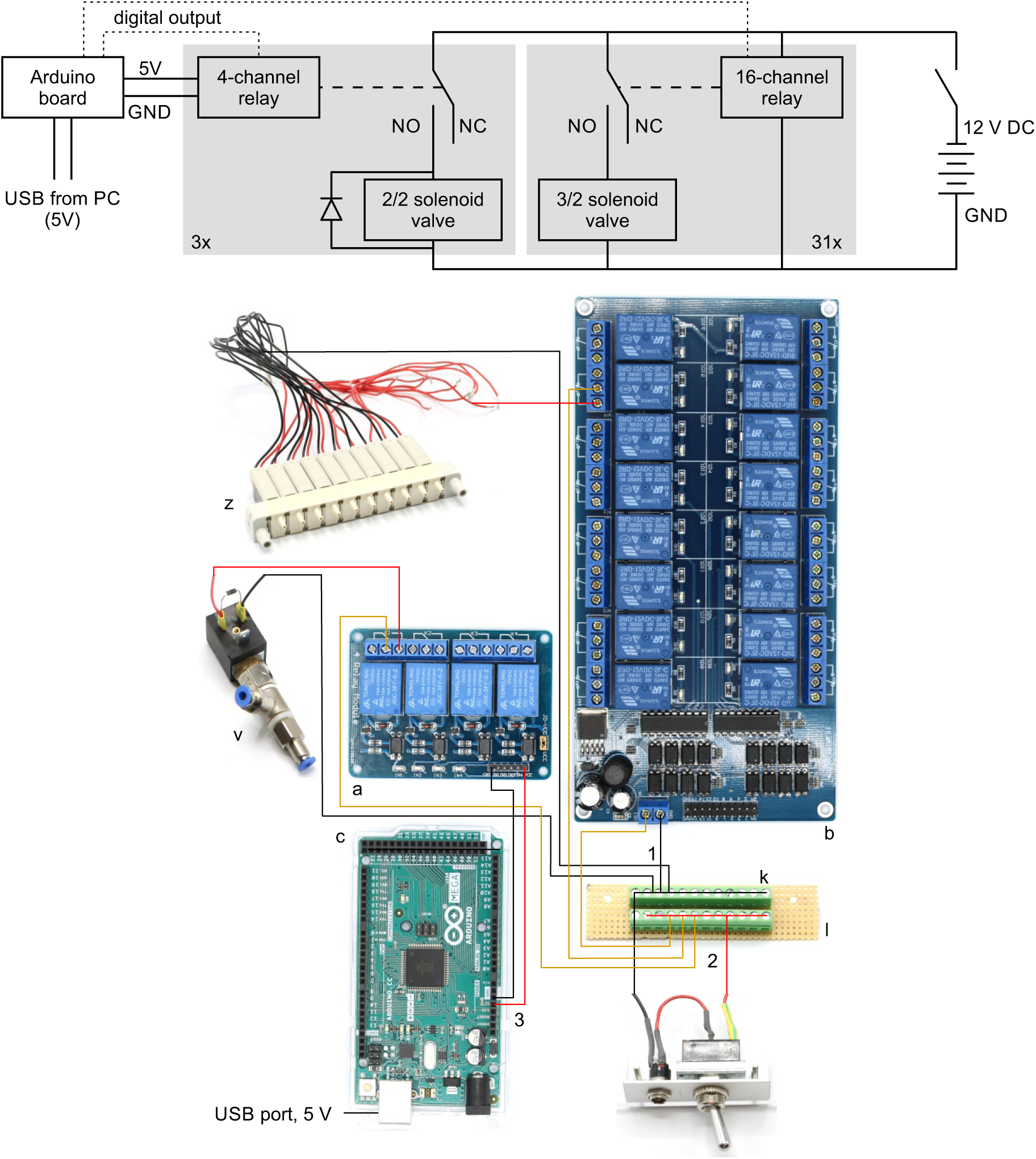

**Top: Electrical circuit wiring diagram for valves, relays and Arduino board.**

This wiring diagram shows that all solenoid valves (z, v) and the two 16-channel relay board (b) are powered through the 12 V DC power supply (80 W, 12 V / 6,67 A). The 4-channel relay is powered through the Arduino board which receives a 5 V power supply from the USB cable. The grey boxes indicate that this circuit is replicated for each valve in parallel. The relays receive digital signals from the Arduino boards (dotted lines) and switch the contacts from NC (normally closed) to NO (normally open) which brings the solenoid valve into a different state. Aflyback diode is necessary for the 2/2 solenoid valves to protect from voltage spikes.

**Bottom: Photographic wiring scheme of circuit.**

1. Ground conductor is connected to all black lines through the soldered screw terminal block on the stripboard (k, l).
2. The 12 V conductor (red) is connected to all yellow lines through the other soldered screw terminal block. These yellow wires carry 12 V to power the 16-channel relay board (b), the individual 3/2 solenoid valves (z) and the 2/2 solenoid valves (v).
3. The 4-channel relay board (a) is powered through the 5 V output pin of the Arduino board (c) which is supplied through the USB-cable from the PC.

#### Assembly guide for pressure control device

0. Prepare the acrylic sheets.

1. Screw in 10 mm metal spacers (h) in M3 threads of the base plate.
2. Place 16-channel relay (b), 4-channel relay (a) and Arduino Mega (c) on the corresponding positions and fix them with the 25 mm metal spacers (i).
3. The Arduino only needs 2× 25 mm spacers to fix it.
4. Connect the input pins of the 16-channel relay with digital outputs of the Arduino (e.g. 22-37) using the jumper wire (d) and document the corresponding pin numbers.
5. Connect the ground pins of the 16-channel relay and the Arduino.
6. Connect the input pins of the 4-channel relay with digital outputs of the Arduino (e.g. 10-13) and document the corresponding pin numbers. Also connect the ground pins.
7. Connect the 5V power pin of the Arduino with the VCC pin of the 4-channel relay to power it (cable not shown in picture).

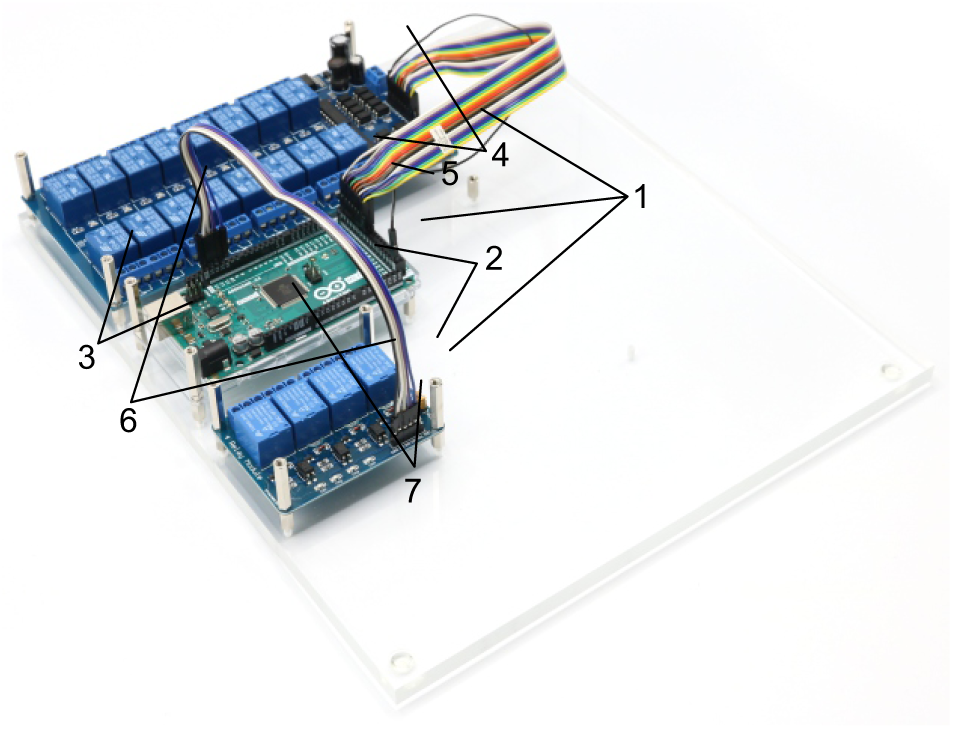
8. Cut 31 yellow isolated copper wires at a length of 15 cm each and strip the ends of the cables.
9. Connect the non-manifold 3/2 solenoid valve (y, S070**C**-6BG-32) to a relay on the 16-channel relay by connecting the red cable to the normally open contact and connecting the yellow cable to the common contact.

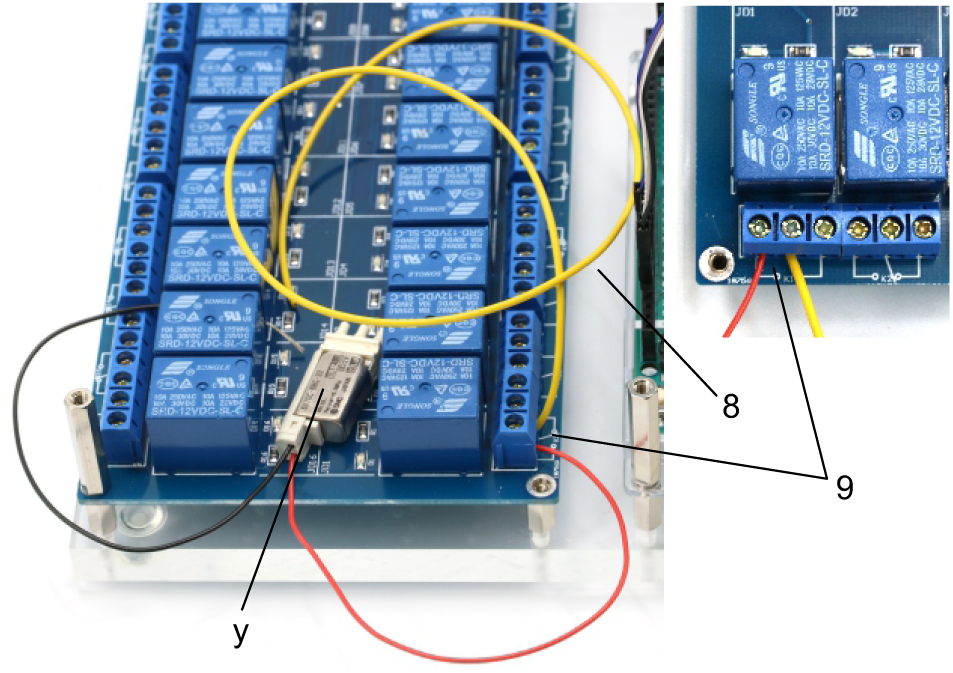
10. Repeat step 9 with all 11 non-manifold 3/2 solenoid valves (y, S070**C**-6BG-32). Document which relay is connected to each solenoid valve. This will be important for the programmed control of each valve in the Matlab code.

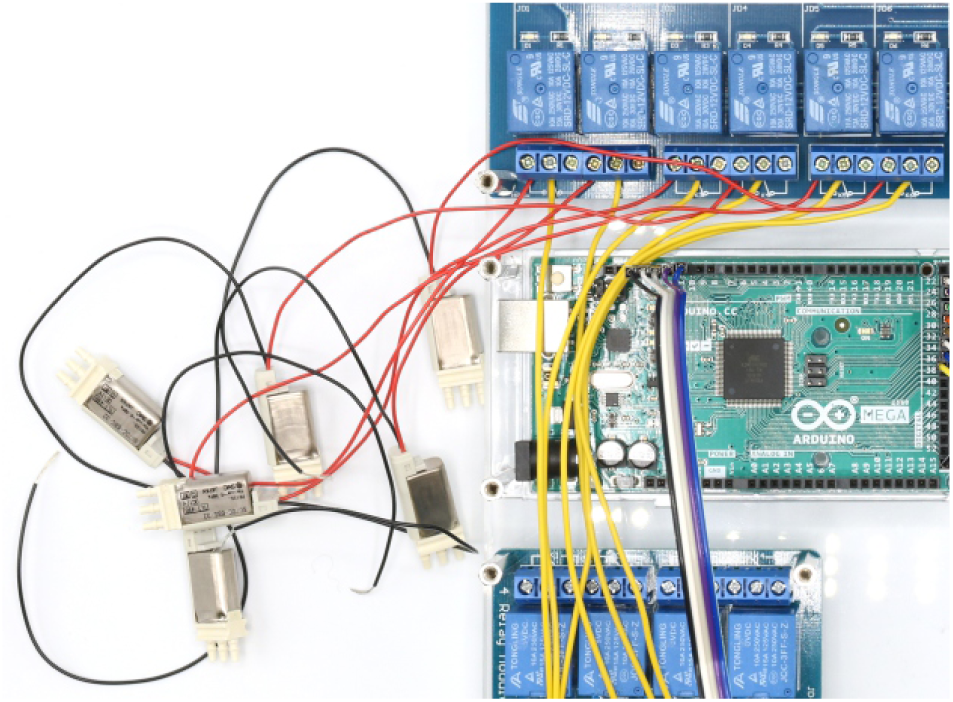
11. Assemble two arrays of 10 manifold 3/2 solenoid valves (z, S070**M**-6BG-32) as described in the product information.

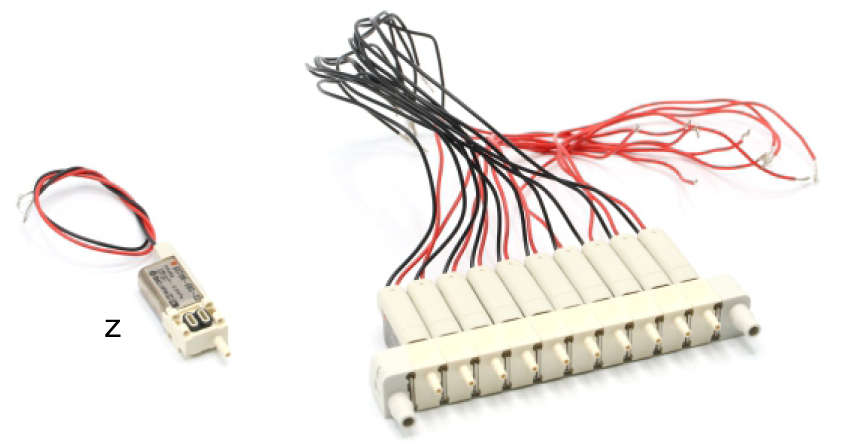
12. Connect 5 manifold 3/2 solenoid valves (z, S070M-6BG-32) to the remaining relays on the 16-channel relay. Connect the red cable to the normally open contact and the yellow cable to the common contact. Document which relay is connected to each solenoid valve.

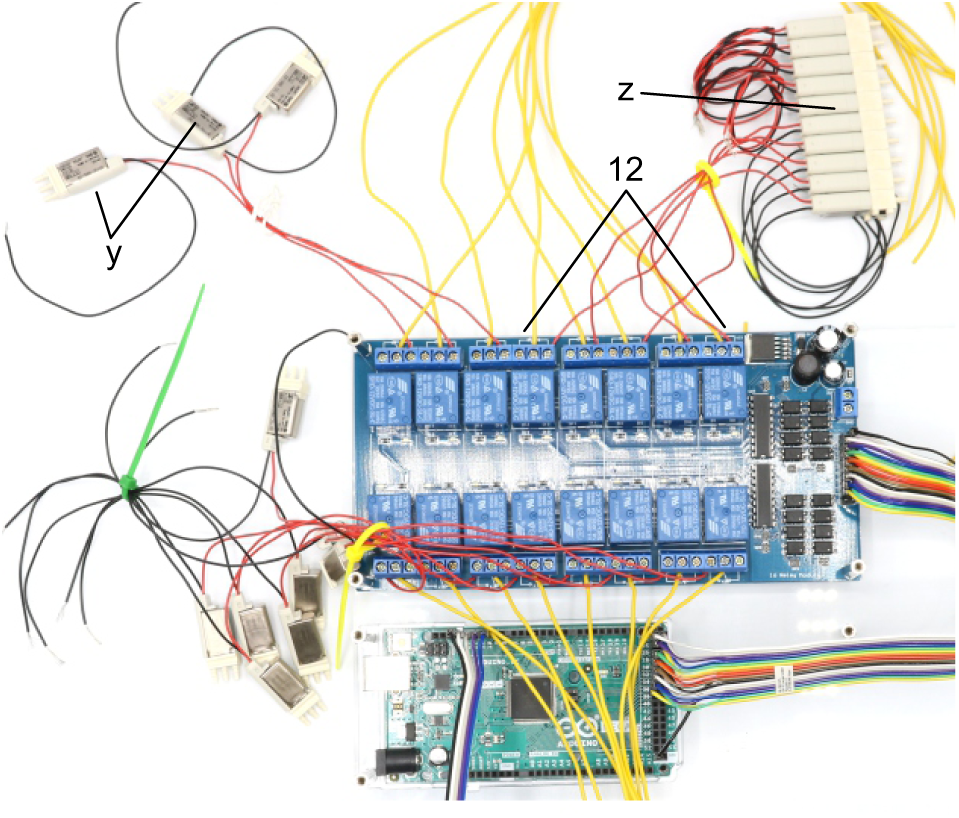
13. Attach the second 16-channel relay (b) on top of the other 16-channel relay and fix it with 25 mm spacers (i)
14. Join remaining 5 manifold 3/2 solenoid valves (z, S070M-6BG-32) of the first assembled array with relays of the top 16-channel relay by connecting the red cable to the normally open contact. Connect a yellow cable to the common contact of every relay. Document which relay is connected to each solenoid valve. Pay special attention to order of connections. Intersections of red cables should be avoided. The valve manifold will be oriented as shown in the picture.

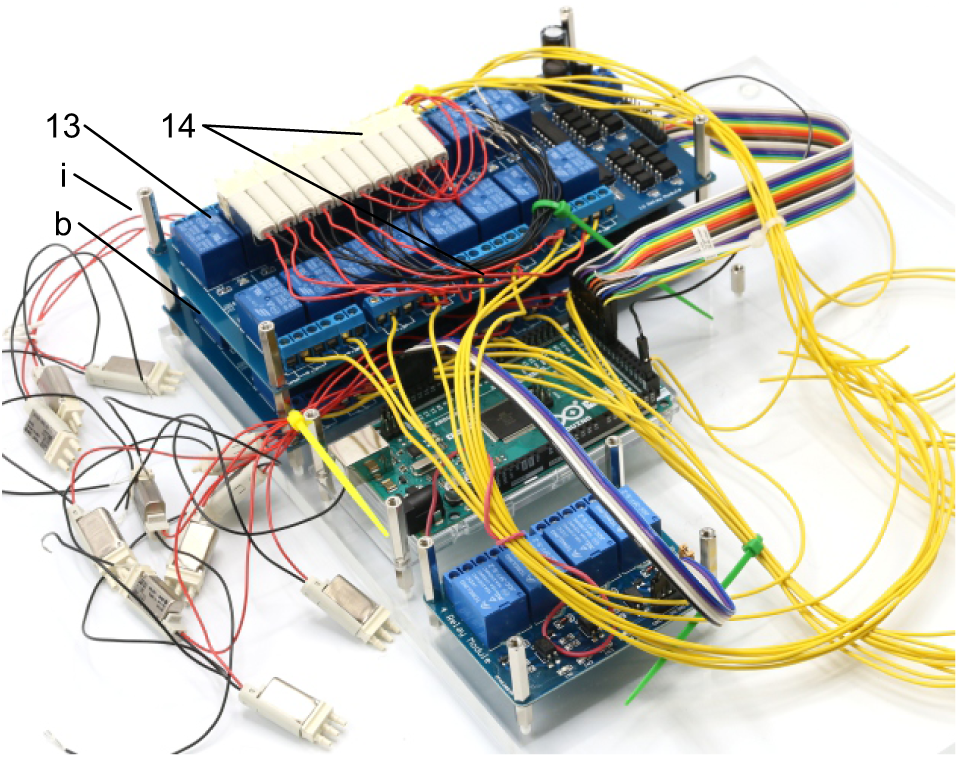
15. Connect the input pins of the top 16-channel relay with digital outputs of the Arduino (e.g. 38-53) using the jumper wire (d) and document the corresponding pin numbers. Connect the ground pins of the 16-channel relay and the Arduino.

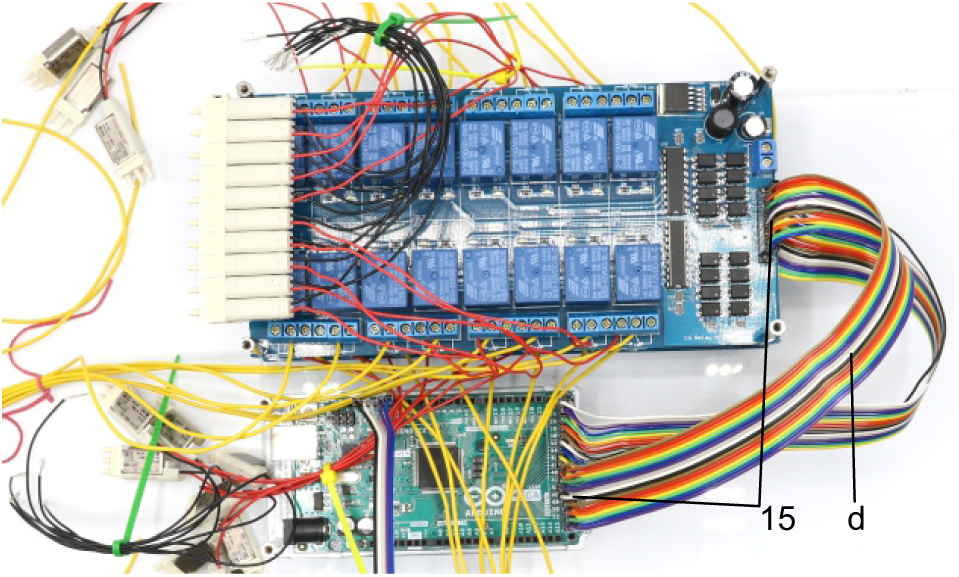
16. Assemble second array of ten manifold 3/2 solenoid valves (z, S070M-6BG-32) and connect these to relays of the top 16-channel relay by connecting the red cable to the normally open contact. Document which relay is connected to each solenoid valve.

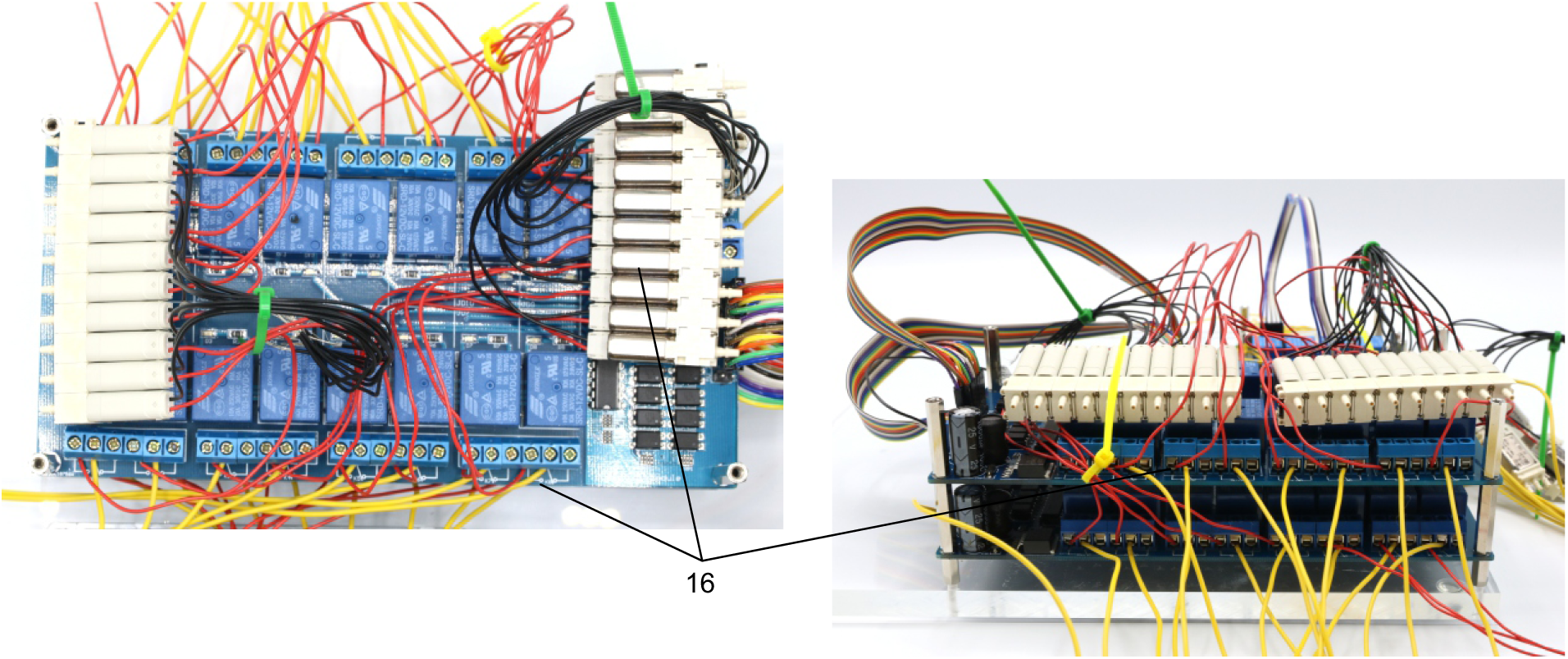
17. Solder red and black wires (approx. 10 cm) to contacts of 3× 2/2 solenoid valves (v).
18. Include a diode (1N4007) between the contacts to avoid current from capacitor discharge damaging the remaining circuit.
19. Connect 2/2 solenoid valves to the 4-channel relay. Connect the red cable to the normally open contact and a yellow cable (approx. 10 cm) to the common contact. Document which relay is connected to each solenoid valve.
20. Screw on push-in fittings (s, q) onto the 2/2 solenoid valves.

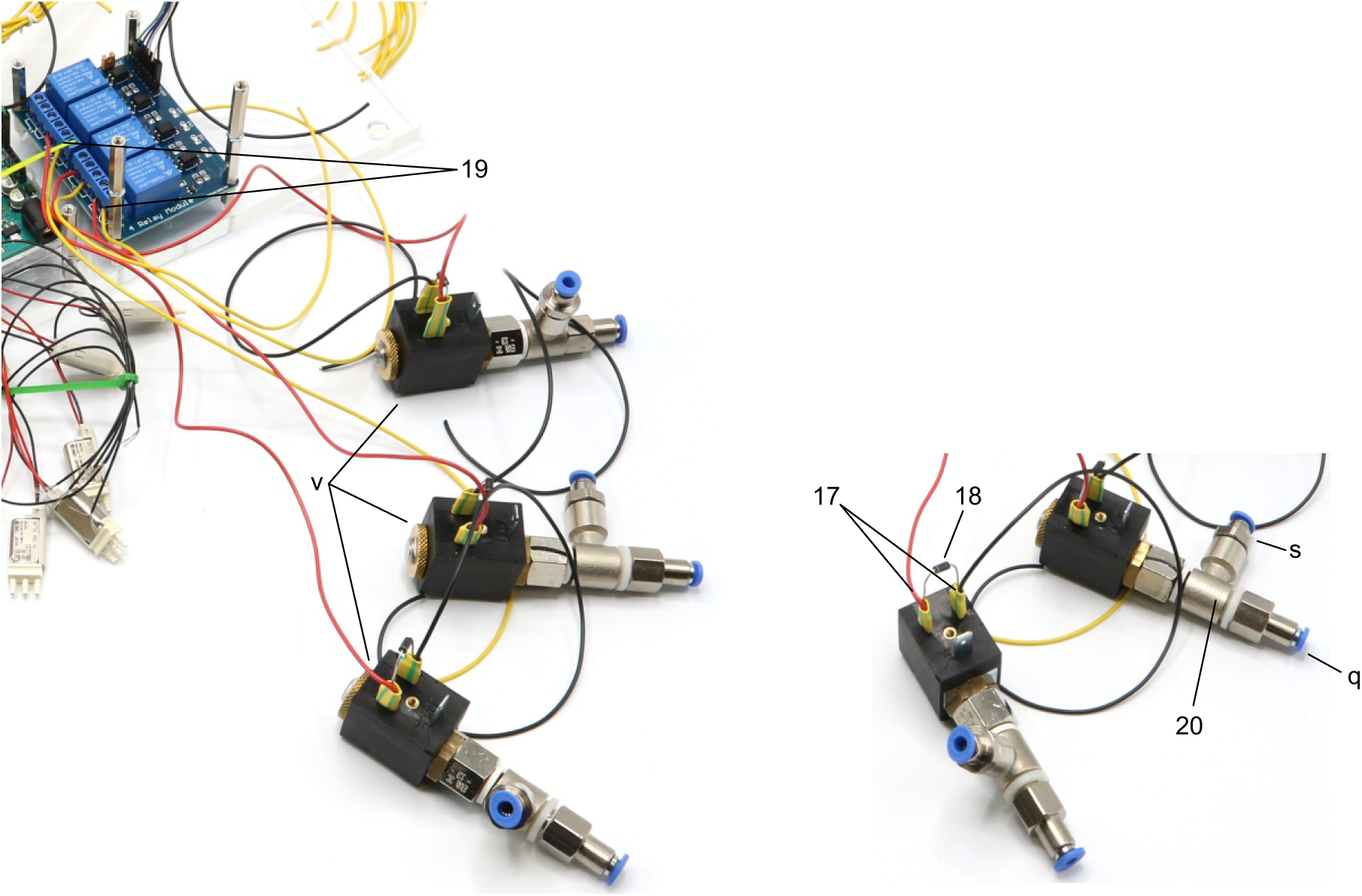
21. Drill holes (3, 8, 12 mm) in aluminium angle (e) as shown on the layout.
22. Attach the toggle switch (f) to the 12 mm hole and the DC barrel power connector (g) to the 8 mm hole.
23. Solder a black wire (approx. 10 cm) to the neutral conductor of the power connector. This wire will connect to all ground wires (black wires) of the valves and relays.
24. Solder a red wire to the DC conductor of the power connector and to a contact of the ON/OFF switch.
25. Solder another red wire (approx. 10 cm) to the other contact of the toggle switch. This wire will condcut 12V for all valves and relays (yellow wires).

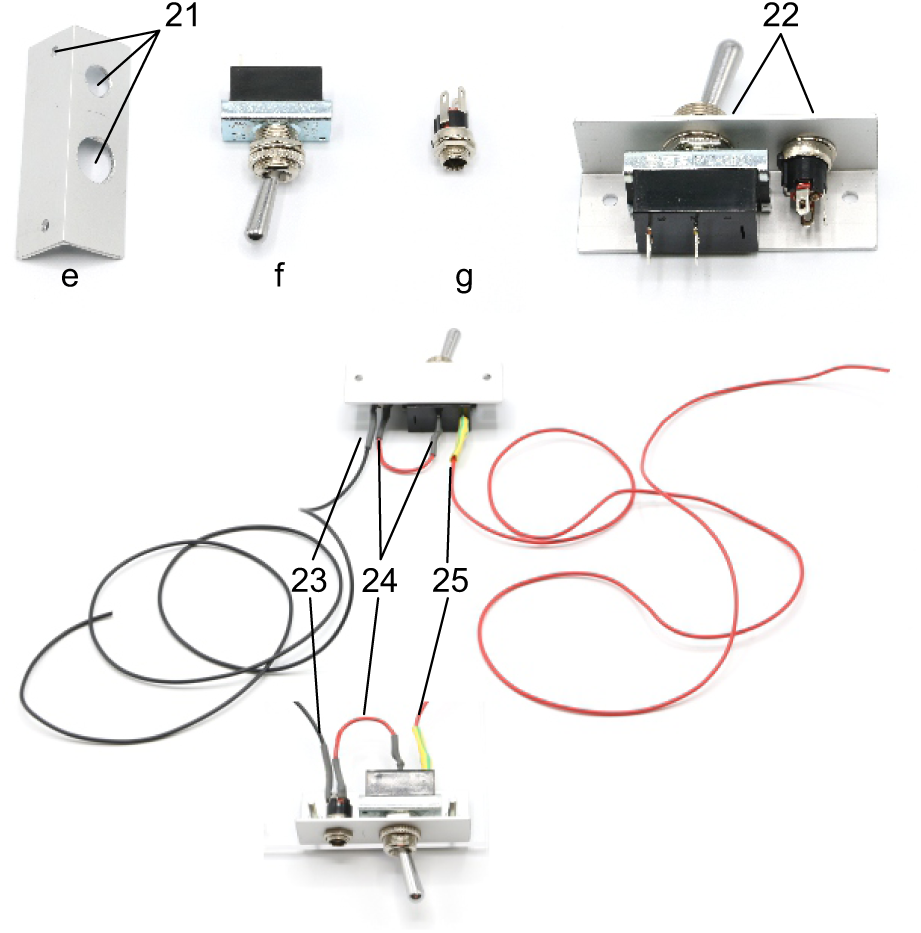
26. Cut the stripboard to a 25×95 mm rectangle with longitudinal orientation of the conducting striples.
27. Drill two 3 mm holes, 82 mm apart and 7 mm away from the edge. This stripboard will be fixed onto the top acrylic plate above the 16-channel relays.
28. Solder both screw terminal blocks (k) onto the stripboard so that all contacts of one block are connected.

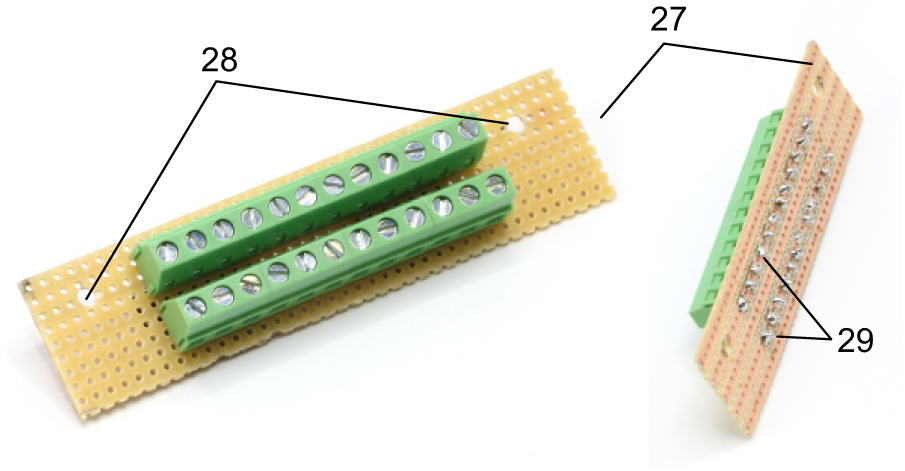
29. Screw in second 25 mm metal spacer (i) on top of all metal spacers to create level pillars of metal spacers for the upper acrylic plate.
30. Mount upper acrylic plate (5 mm) on top of the 25 mm metal spacers (i).
31. Use 20 mm M2 bolts to attach the non-manifold 3/2 solenoid valves (y) at the location of the 2 mm holes on the upper acrylic plate. Use M2 nuts below the acrylic plate to fix the valves.

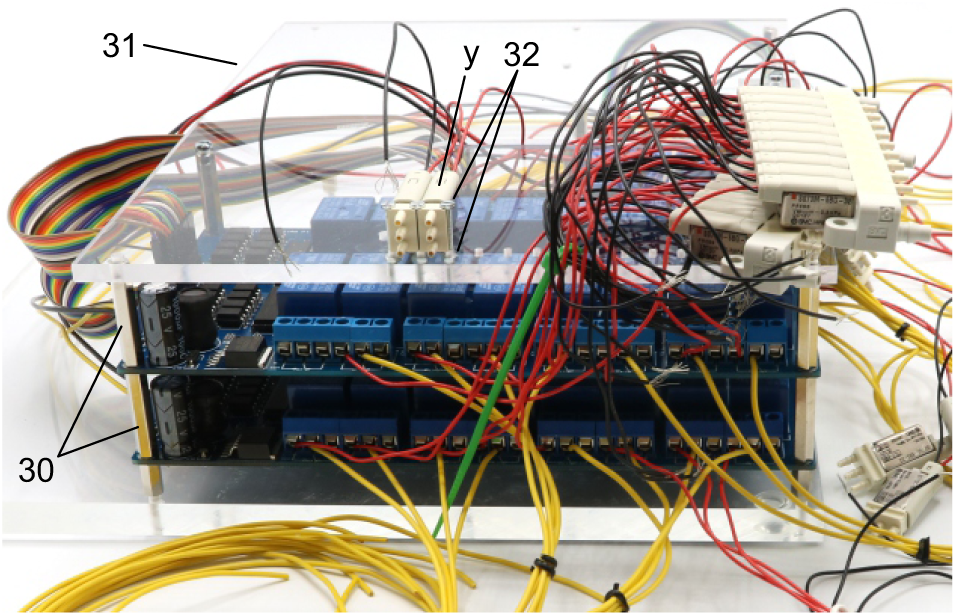
32. Repeat step 32 for 10 of the 11 non-manifold 3/2 solenoid valves (z). Document position of each valve, their respective relay and connected Arduino digital output pin.
33. Screw in two 25 mm metal spacers (i) next to the first level of non-manifold solenoid valves.
34. Attach the other two solenoid valve manifolds (z) on top of the metal spacers separated by 20 mm plastic spacers (j) and fixed using 35 mm M3 bolts. Pay attention to avoid twisting of the cables. Route the cables either through the notch or around the upper plate.
35. The 11. non-manifold solenoid valve is later attached on the top plate.

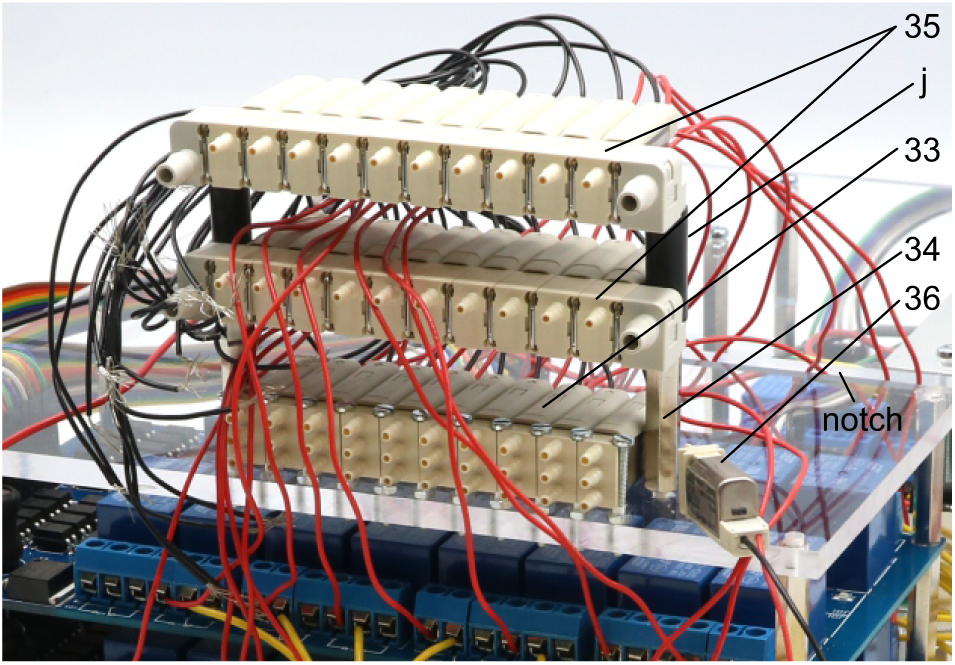
36. Attach vacuum ejector (t), equipped with G1/4 push-in fittings (r) and silencer (u), and three 2/2 solenoid valves (v), equipped with G1/8 push-in fittings (s, q), onto the top plate using double-sided adhesive tape.
37. Use 25 mm M3 bolt to attach the push-in Y-fitting (o).
38. Use 35 mm M3 bolt to attach the multiple distributor with 4 outlets (m, p).
39. Attach the angle with toggle switch and DC power connector (step 21-25) underneath the top acrylic plate, above the Arduino and use M3 bolts to fix it.
40. Attach the cut stripboard with soldered screw terminal blocks (step 27-29) to the top plate using two 25 mm M3 bolts and two 10 mm plastic spacers (j).
41. Connect all yellow wires from the relays and the 16-channel relay boards to one screw terminal block which also connects the red 12V phase conductor from the toggle switch (step 25). This distributes the 12V power from the DC power connector to the relays and thereby to the solenoid valves through the red wires.
42. Connect all black neutral wires from the solenoid valves and the 16-channel relay boards to the other screw terminal block which is also connected with the black neutral conductor wire from the DC power connector (step 23).
43. Use double-sided adhesive tape to attach the 11. non-manifold solenoid valve (step 36) onto the top plate, adjacent to other 3/2 solenoid valves.
44. Use 10 mm M3 bolts on remaining holes.

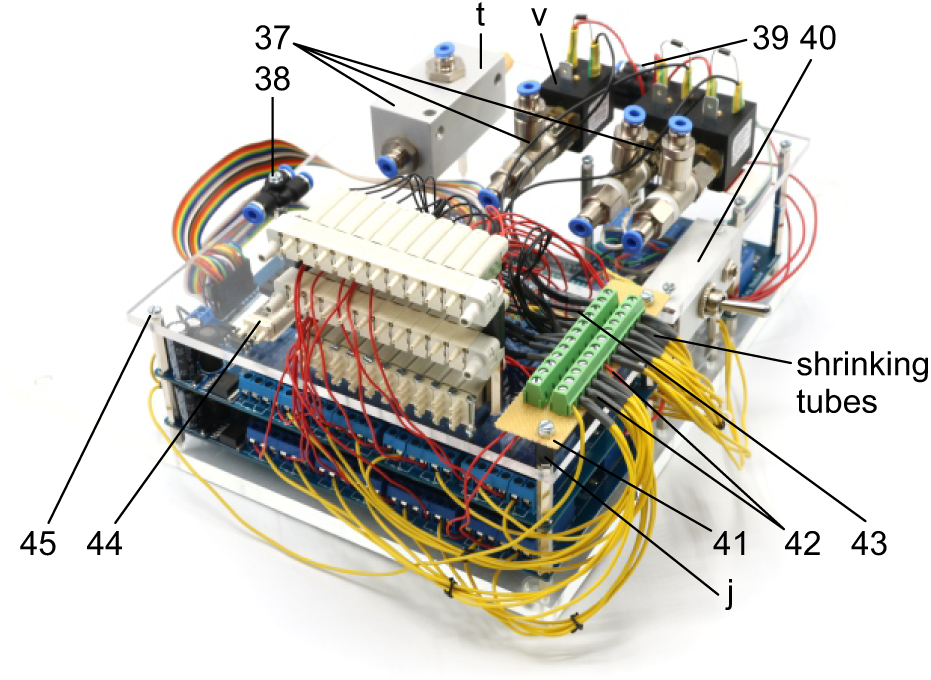

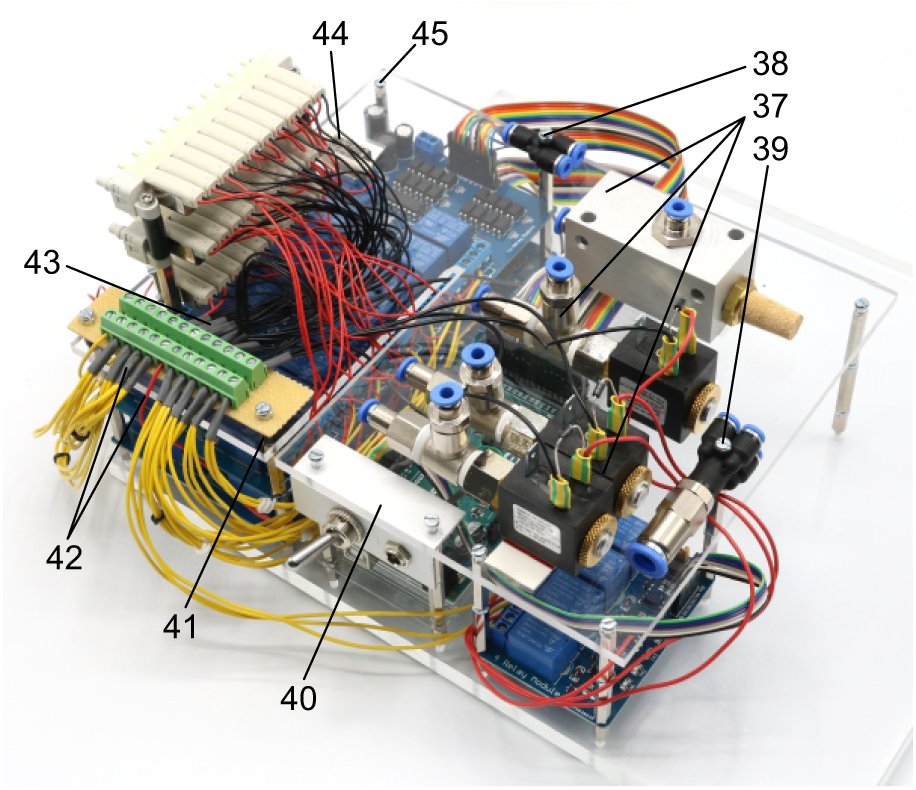 The following steps connects the 3/2 valves to enable application of different pressures to individual pipette holders. For schematic depiction, please see respective figure.
45. Cut 10 silicon tubes (diameter 2/4 mm) with a length of 9.5 cm, 10 silicon tubes with a length of 6 cm and 10 silicon tubes with a length of 8 cm.
46. Connect one end of the 9.5 cm tubes to the 2A nozzle of the top level valves and the other end to the lower 3R nozzle of the respective first level valves.
47. Connect one end of the 6 cm silicon tubes to the 2A nozzle of the second level valves and the other end to the middle 1P nozzle of the respective first level valves.
48. Connect one end of the 8 cm tubes to the top 2A outlets of the first level valves. The other end can be connected to tubes that directly connect to the respective pipette holders.

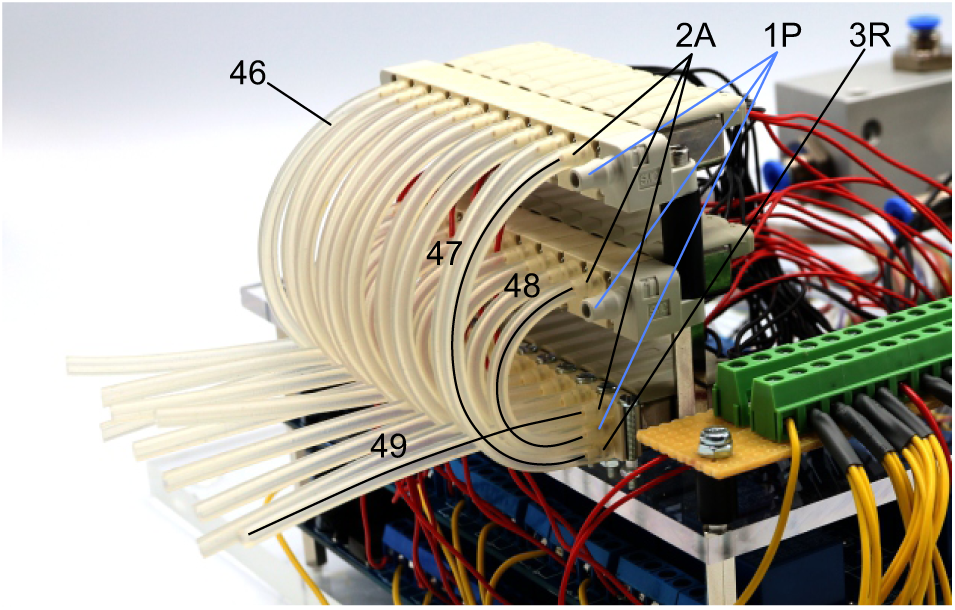 The following steps connects the 2/2 valves and vacuum ejector to the 3/2 valve manifolds to generate the necessary pressures for the cleaning procedure. The tubes highlighted in red transfer the 1 bar positive pressure. The tubes highlighted in yellow transfer the negative pressure for suction. Arrowheads indicate direction of air flow.
49. Connect the PE-tubes (outer diameter 4 mm, inner diameter 2 mm) as shown between the 2/2 valves and the vacuum ejector. Connect these to the Y-fitting.
50. Connect the Y-fitting to the 3R nozzle of the second level 3/2 valve manifold through a silicone tube (diameter 4/6 mm).

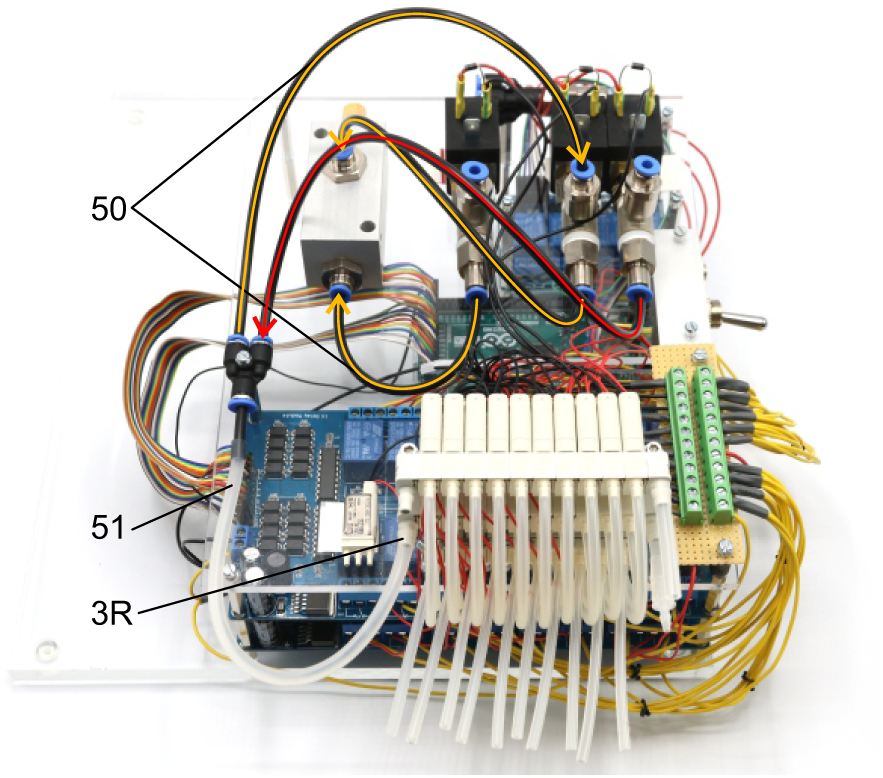 The following steps connect the 1-10 bar pressure regulators (w) to the pressurized air supply and the 2/2 solenoid valves.
51. Position the pressure regulators (w, x) on the base plate and fix them using double sided adhesive tape. We found this to be of sufficient stability. Pay special attention to the intended direction of air flow depicted on the regulators.
52. Connect the PE-tubes to the multiple distributor with 4 outlets and to the 1-10 bar pressure regulators (w). Then connect those to the 2/2 solenoid valves as depicted. Meaning of colour code and arrowheads correspond to the previous steps.

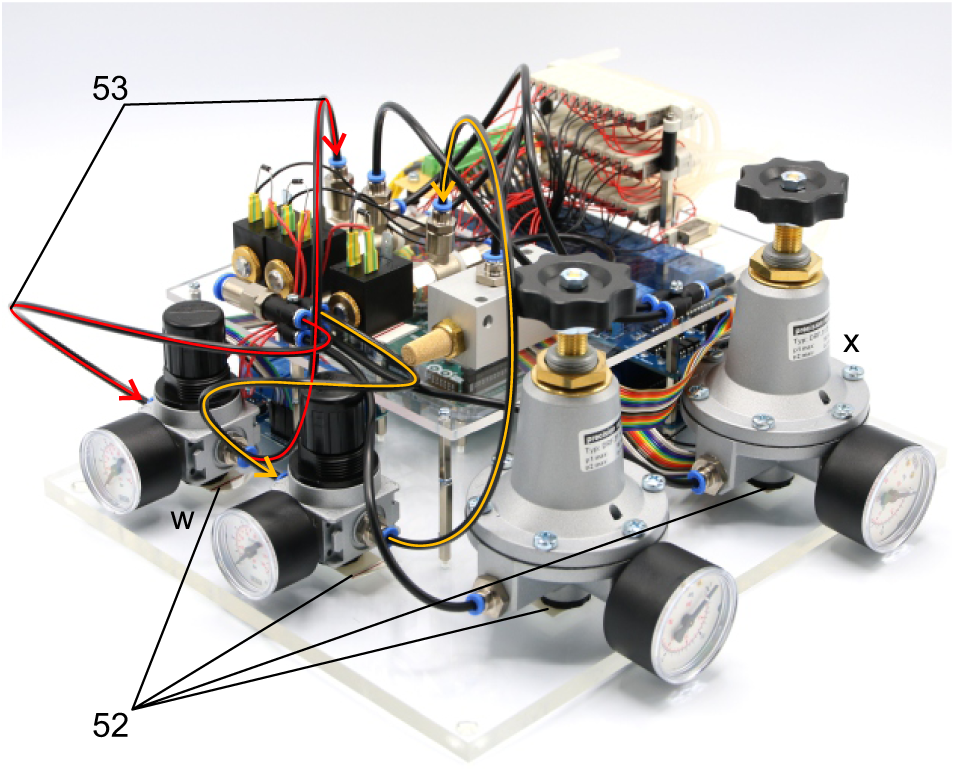 The following steps connect the precision pressure regulators (x, 10-1000 mbar) to the valve manifold to generate HIGH pressure (70 mbar, green lines) and LOW pressure (20 mbar, blue lines).
53. Connect 2 PE-tubes to the multiple distributor with 4 outlets and each to one of the precision regulators. Pay spetial attention to intended direction of airflow on the regulators. Arrowheads indicate direction of air flow.
54. Connect the HIGH pressure regulator (set to 70 mbar) to the middle nozzle of the single 3/2 solenoid valve which is attached to the acrylic plate (green line). Connect the PE-tubing with a 2/4 mm silicone tube using a mini tubing connector(*).
55. Connect the LOW pressure regulator (set to 20mbar) to the 1P nozzle of the top level 3/2 valve manifold. Use a 4/6 mm silicone tube to connect the PE-tubing with the nozzle.

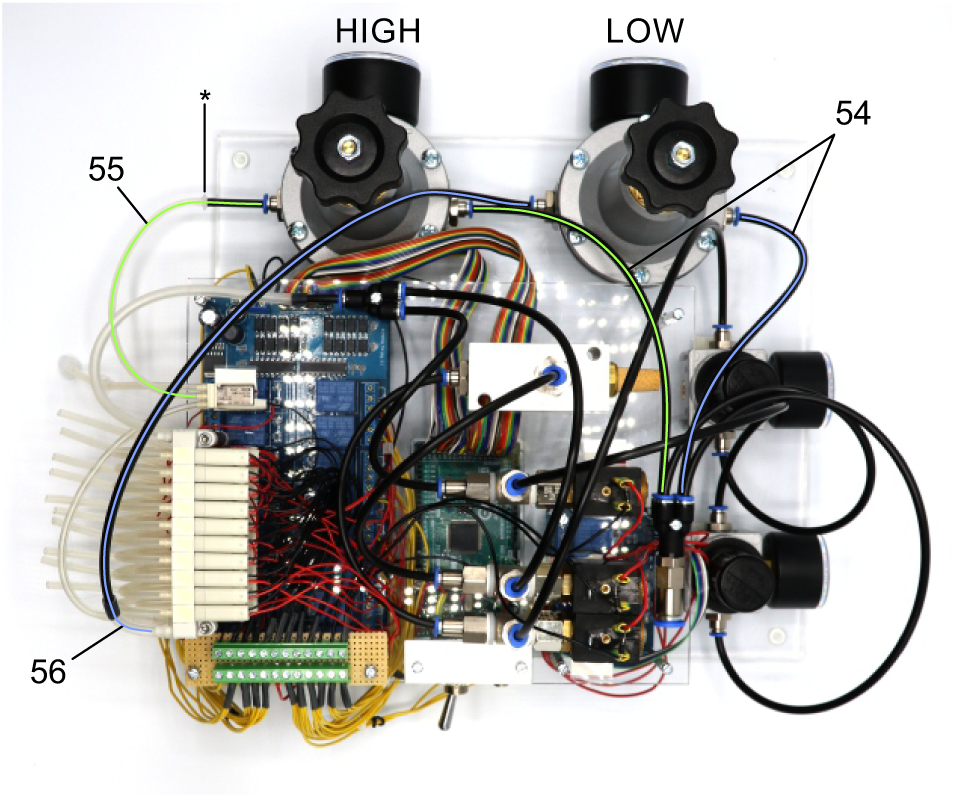
56. Connect the 1P nozzle of the second level solenoid valve manifold to the 3R nozzle of the single 3/2 solenoid valve. Use a 4/6 mm silicone tube to connect the 1P nozzle of the manifold. Connect it with a 2/4 mm silicone tube using a mini tubing reducer.
57. Connect a 2/4 mm silicone tube to the 2A nozzle of the single 3/2 solenoid valve which can be connected to a mouth piece or a syringe for applying pressure during membrane sealing and breakthrough.

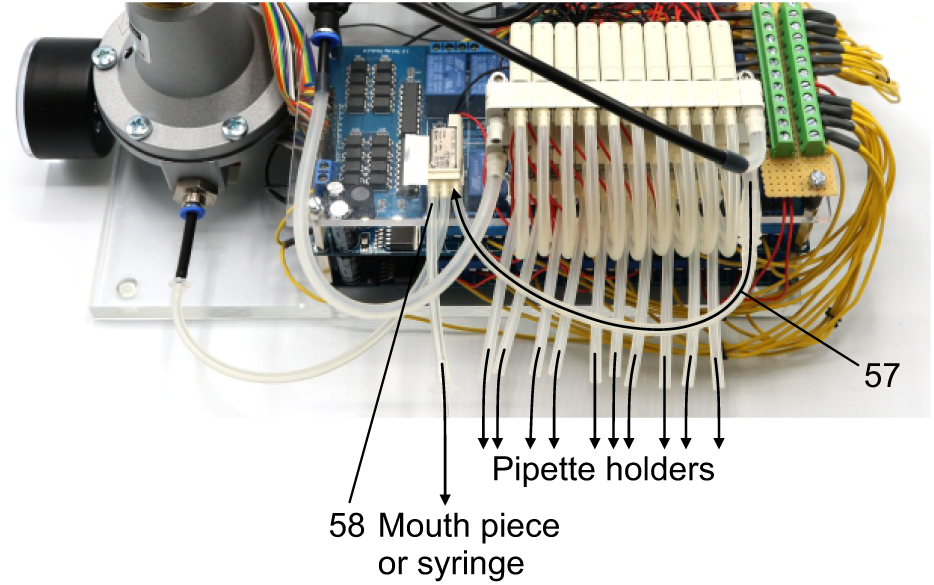
58. Plug in the power supply (12 V / 6.67 A, 80 W) and the USB-cable to the Arduino board.
59. Connect to presurrised air supply using a PE-tubing with 8 mm outer diameter. The air supply should be set at sufficient high pressures (e.g. 5 bar)

Adjust the regulators manually to the desired pressures.

*The pressure regulator upstream of the vacuum ejector needs to be adjusted while measuring the negative pressure generated by the vacuum ejector. It should be set at a pressure at which -350 mbar can be applied to the valve manifold (approx. 3-5 bar).

To control the device, you can use the Matlab code and GUI we provided. One could also control the Arduino board using any custom script. Before the first use, please transfer the mapping of the individual valves onto their relay number and respective Arduino digital output pin into the script (see guide to Matlab GUI).

Now the pressure control device is fully operational.

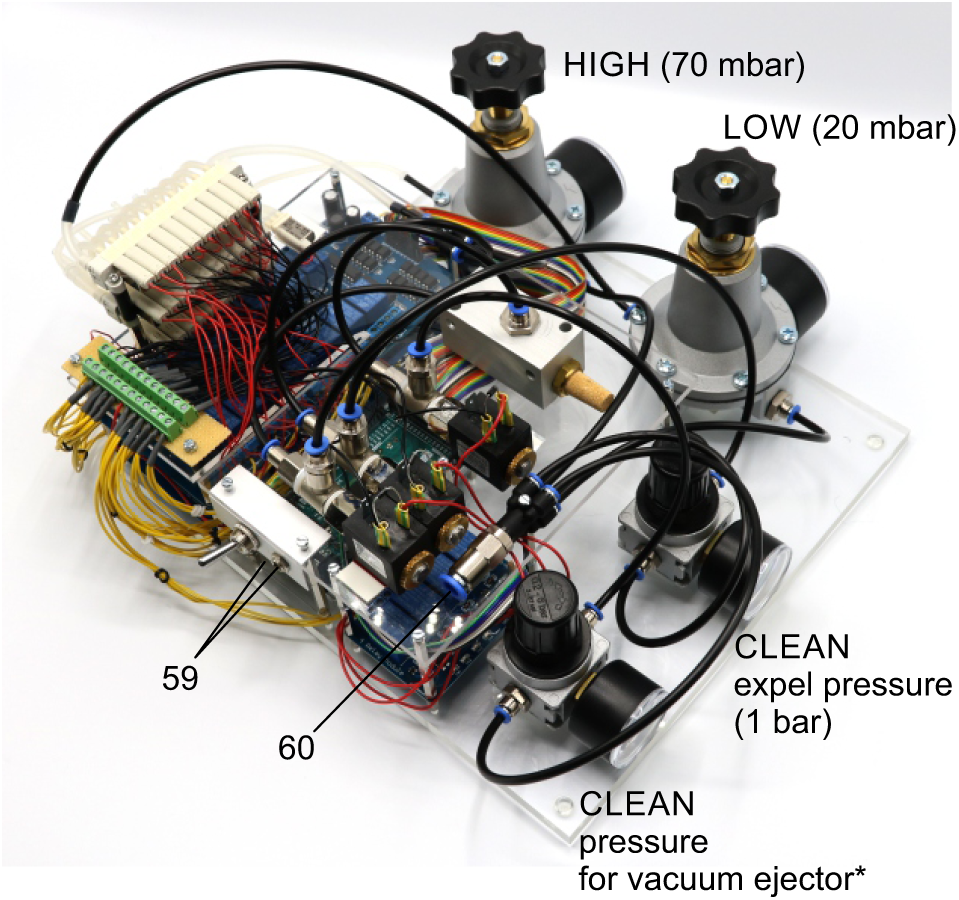

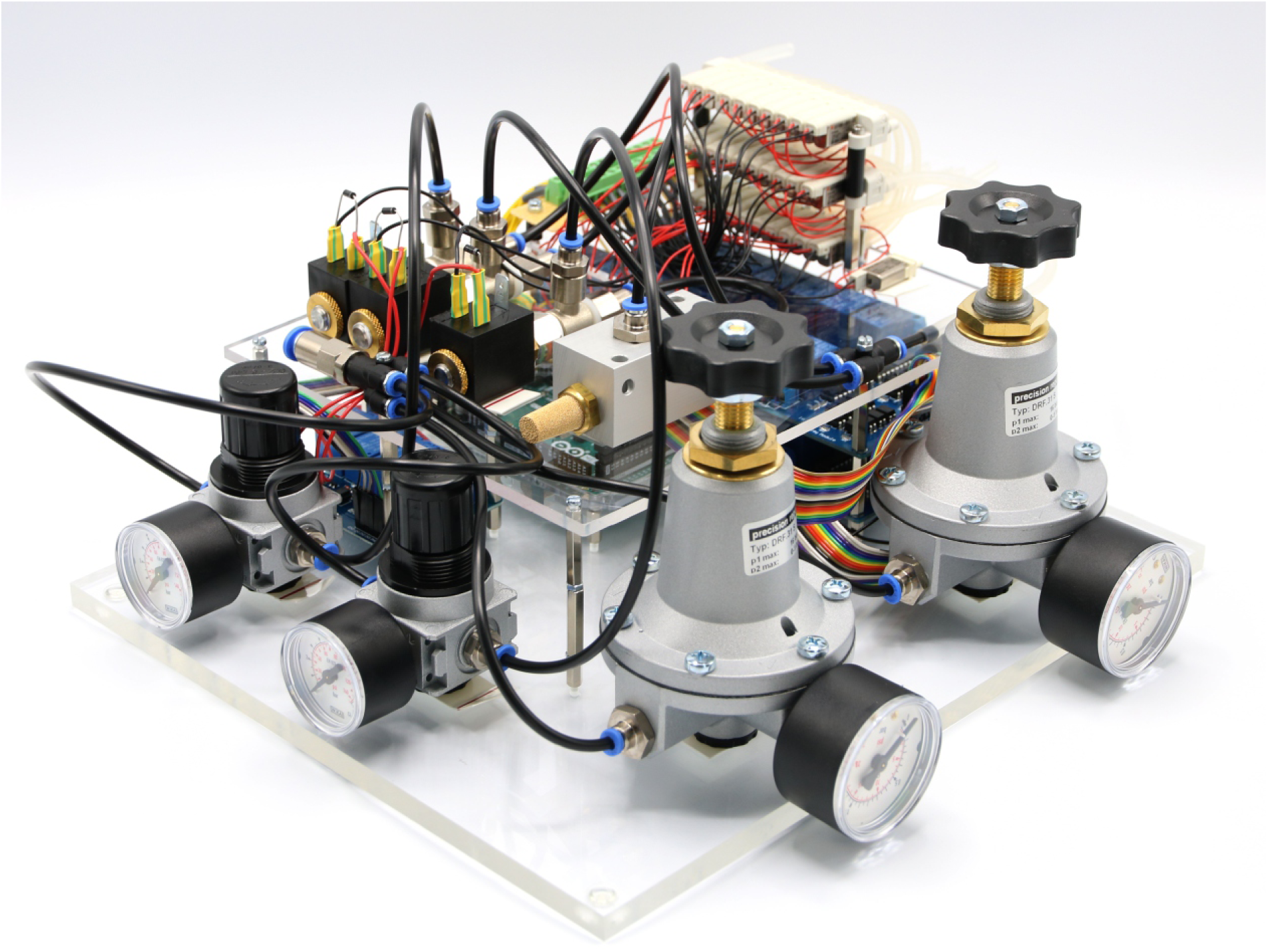

### Matlab code

#### GUI instructions

We developed a graphical user interface to control the pressure system and the automated pipette movements. The Matlab code for the pressure system can be downloaded from our Github repository: https://github.com/neurocharite/multipatch

For further elaborations regarding the movement algorithms, please see the readme file.

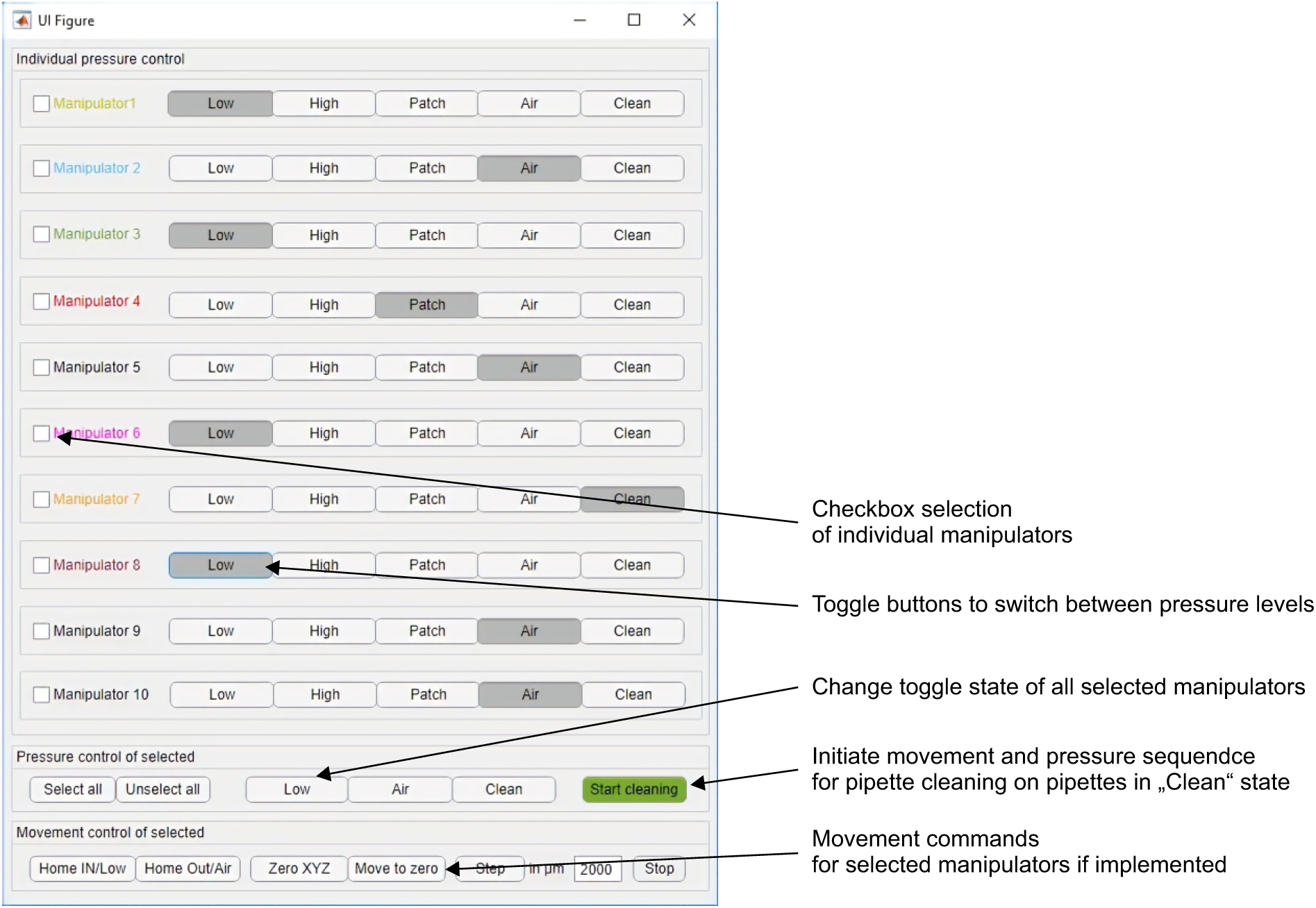

#### Pipette positioning algorithm - principle of operation

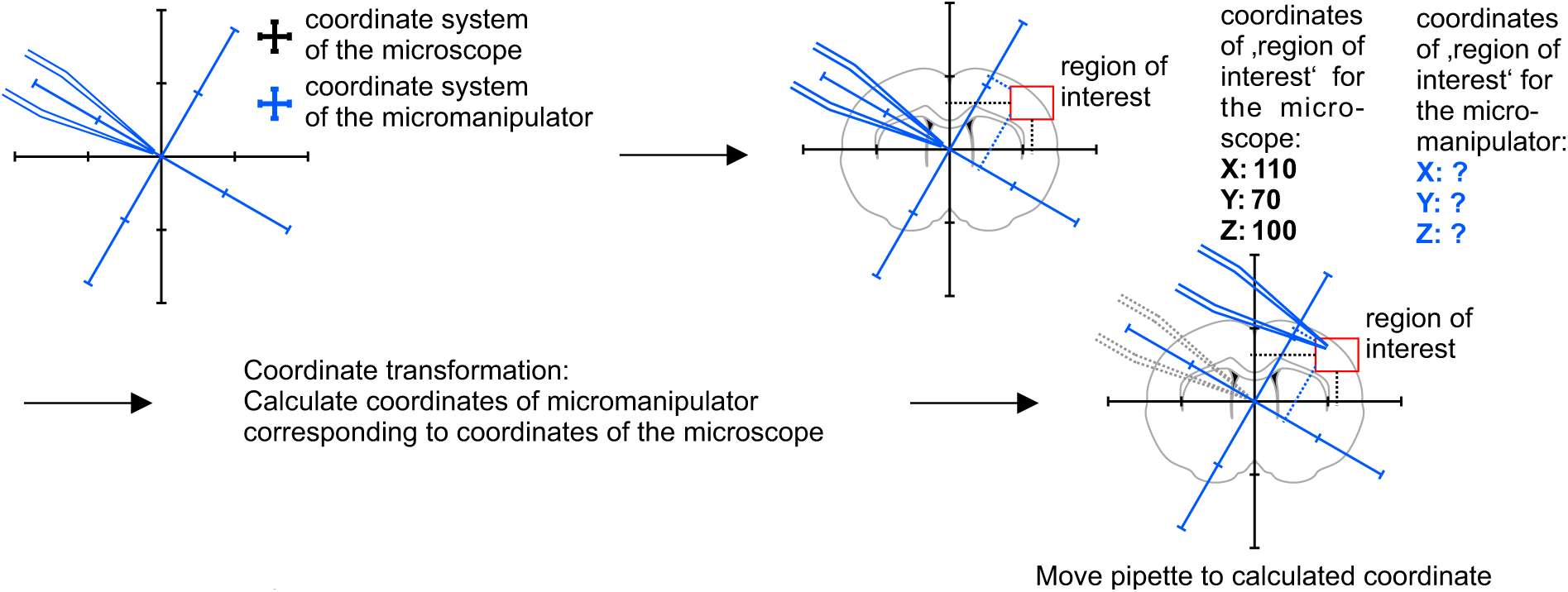

#### Application of the algorithm

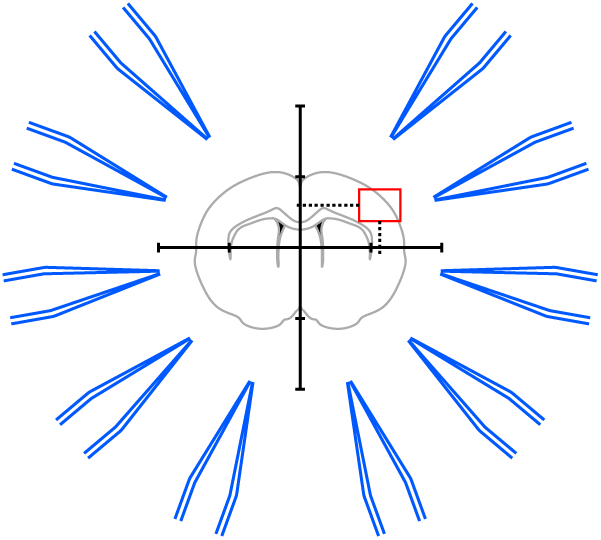

Step 1:

A brain slice is placed under the microscope. The, region of interest’ is located in the coordinate system of the microscope.

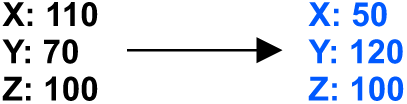

Step 2:

The coordinates of the, region of interest’ are fed into the pipette finding algorithm. The coordinates for each manipulator are computed.

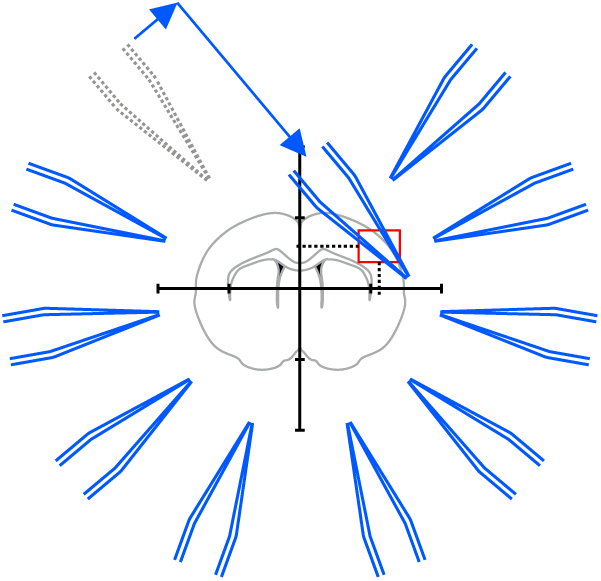

Step 3:

Pipette 1 is moved to the calculated coordinate. Manual adjustments are necessary to due variations in pipette shape.

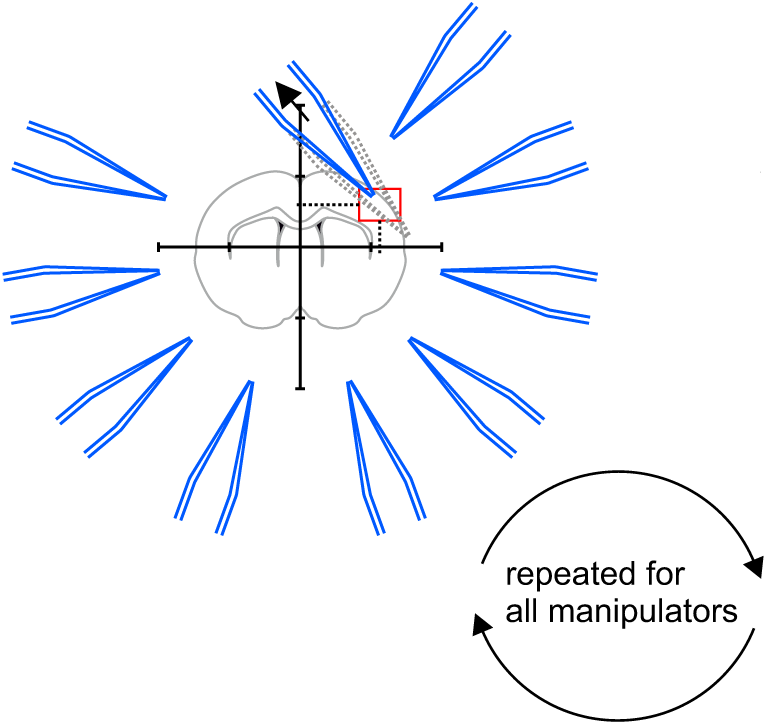

Step 4:

Pipettes should be moved to a designated corner that will prevent collisions with the other pipettes.This position is then saved.

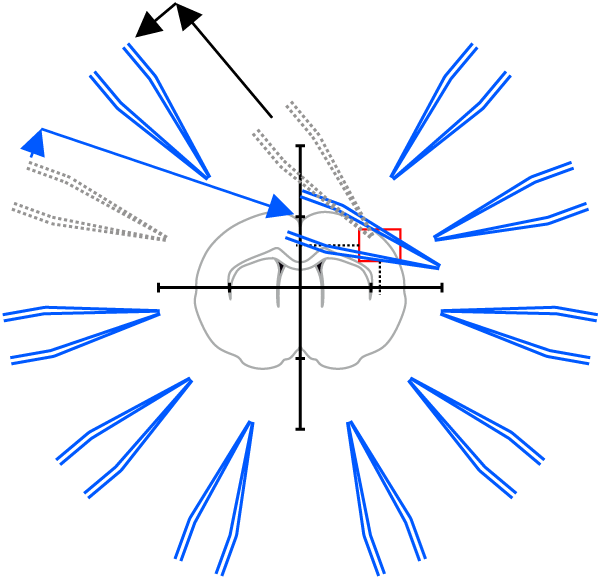

Step 5:

The pipette is moved to its starting position while the next pipette is moved to the position calculated by the pipette finding algorithm.

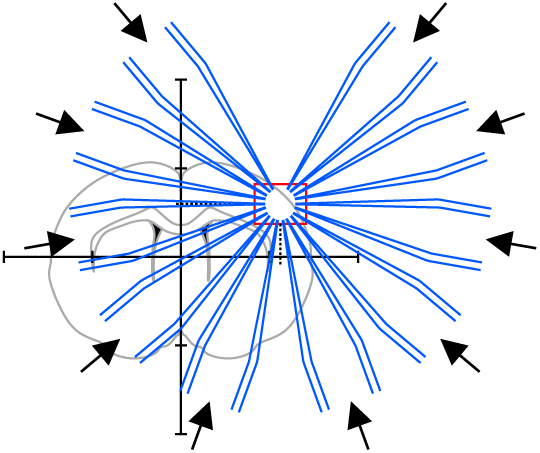

Step 6:

All pipettes are moved to their target positions simultaneously. All pipttes are now positioned in the region of interest

Manipulator coordinates were matched to the microscope coordinates using a rotation matrix and anchored to a common reference point. Whenever a new region of interest is determined in the microscope, these coordinates are then entered into the pipette positioning program and the pipette tip is moved into the new field of view by calculating the offset. To prevent collision with the slice, we performed this positioning process approximately 2500 µm above the slice. Small manual adjustments are frequently necessary due to varying length of the pipettes. After the pipette tip is manually moved to its dedicated section of the field of view, this new “IN” position is saved, and the pipette is moved to its initial “OUT” position. Then the next pipette is moved to the calculated target position and the procedure is repeated until all pipettes were assigned new “IN” positions. At the end, all pipettes are moved to their assigned “IN” positions and lowered in the z-axis to roughly 250 µm above the slice surface to prevent effects of the intracellular solution on the slice. Now, individual whole-cell patch-clamp recordings can be established consecutively. This semi-automated approach reduces the risk of breaking the pipette tip and the time needed to complete pipette positioning to 7-9 minutes.

### Construction drawings

#### Recording chamber for Scientifica manipulator

all measurements in mm dotted lines = edges on underside

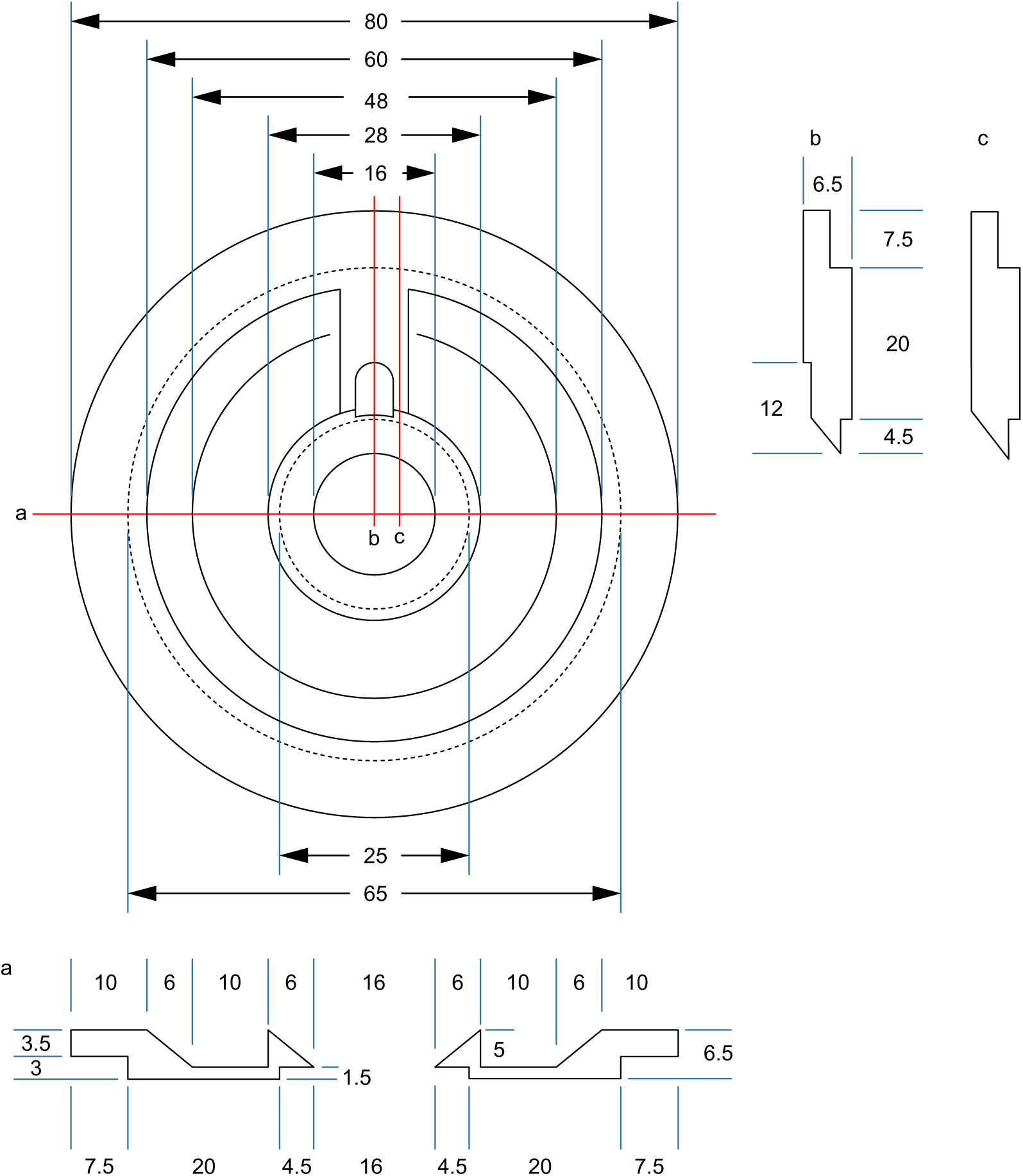

#### Recording chamber for Sensapex manipulator

all measurements in mm

dotted lines = edges on underside

Adjustments are needed for Sensapex micromanipulators. Due to a range of motion in the axial axis of 20 mm, the real horizontal range of motion is less than 20 mm. Therefore the central recording well has a smaller diameter.

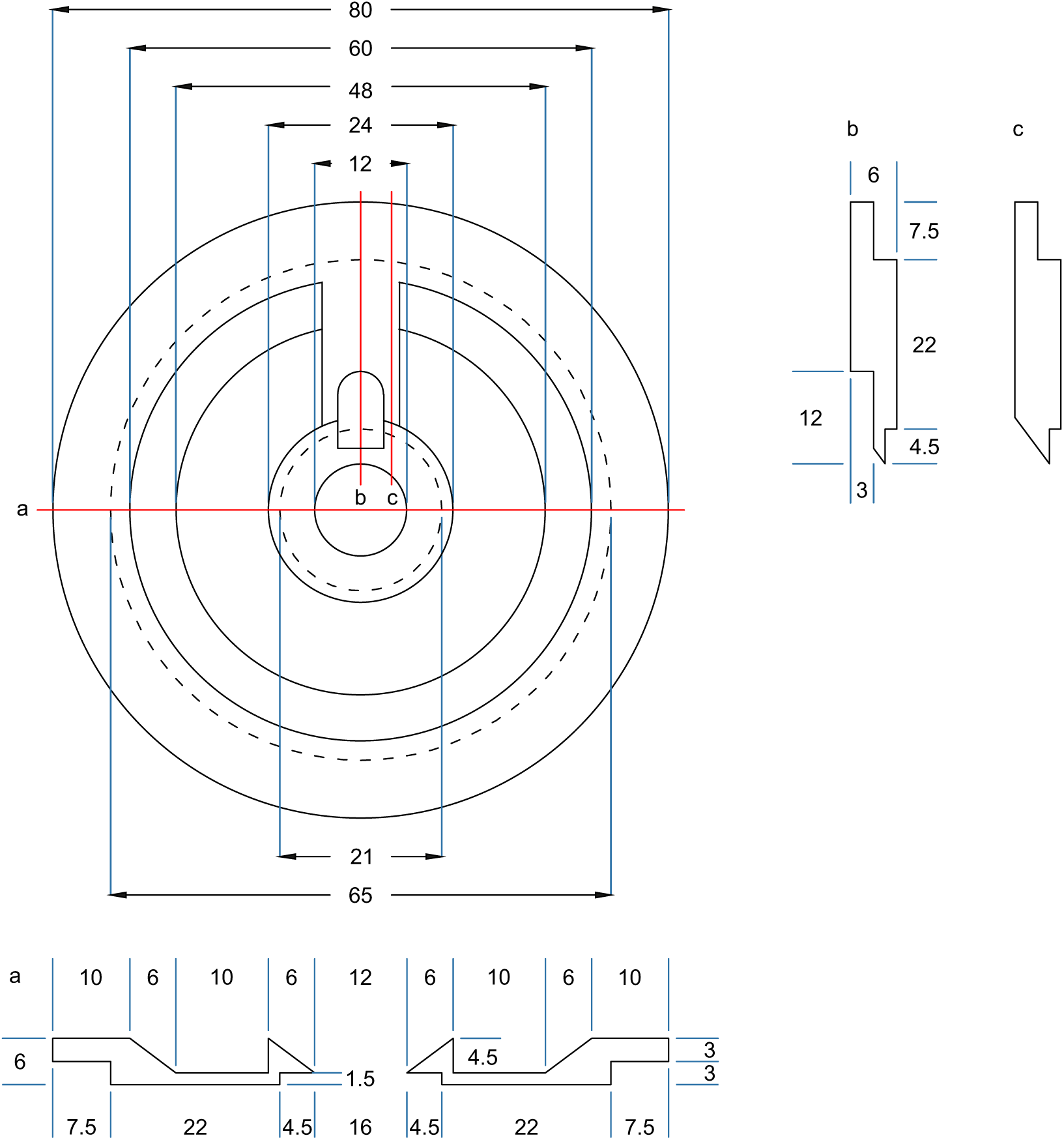

#### Custom stage for Sensapex micromanipulator with Nikon Eclipse microscope

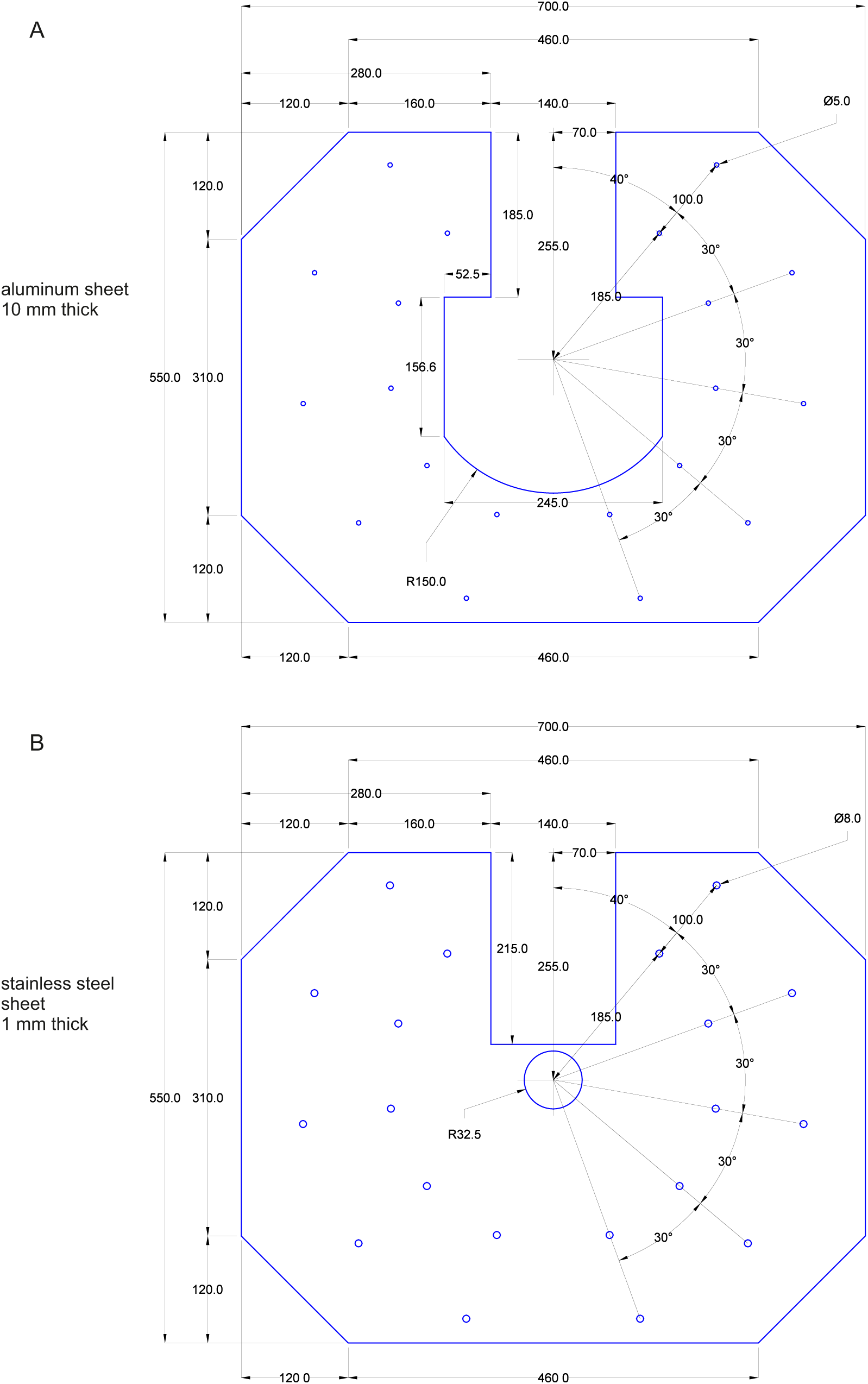

**Supp. Fig., Stage construction plan‘. A** 10 mm thick sheet of aluminum. The sheet is shown in blue. Measurements are shown in black. All measurements are in milimeters. All drilling-holes are 5 mm in diameter (for M6 ISO metric screw threads). **B** 1 mm thick sheet of stainless steel. The sheet is shown in blue. Measurements are shown in black. All measurements are in milimeters. All holes in the stainless steel sheet are 8 mm in diameter. The two sheets have to be glued together before being mounted into the setup with the stainless steel sheet facing upwards.

#### Multi-pipette filling device

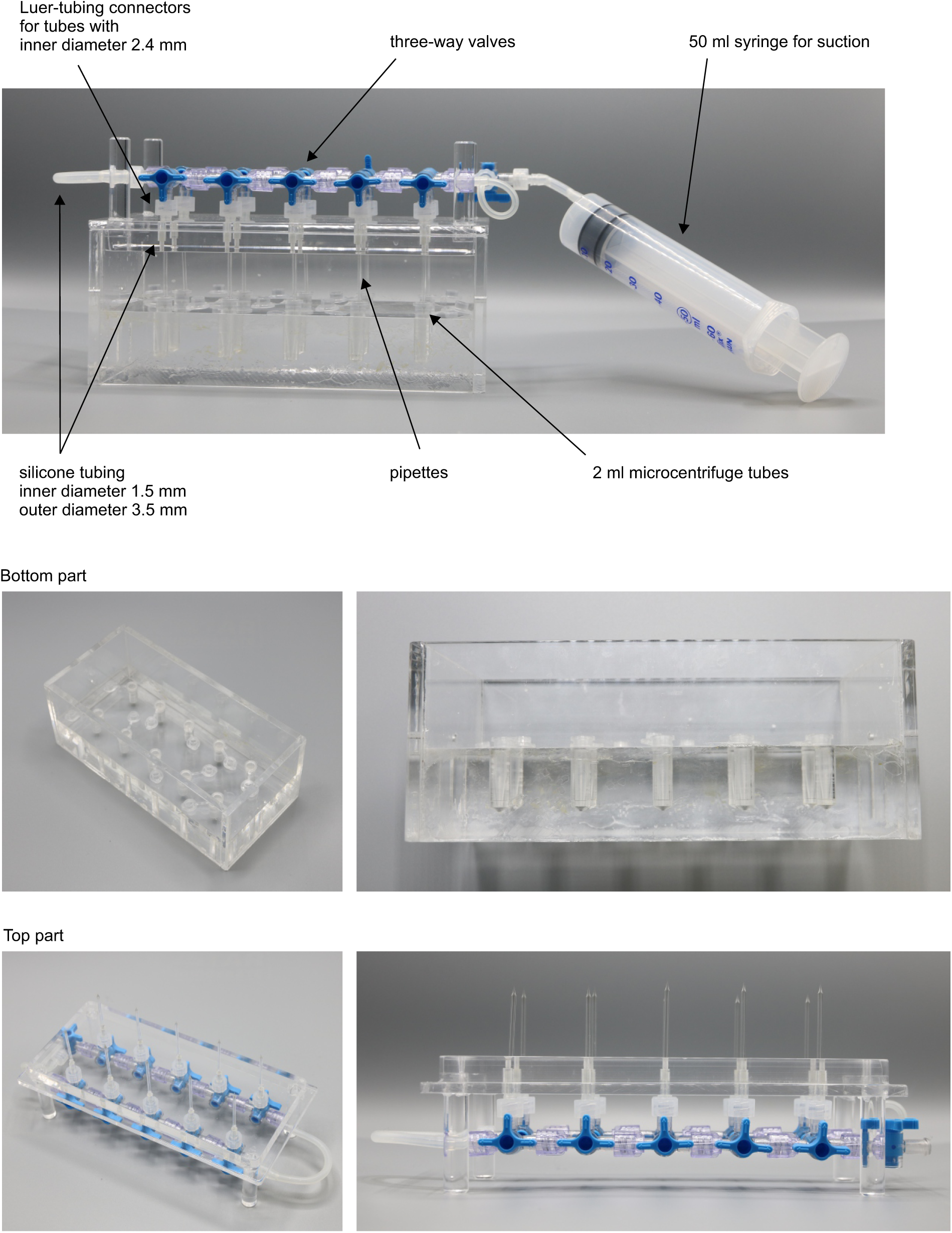

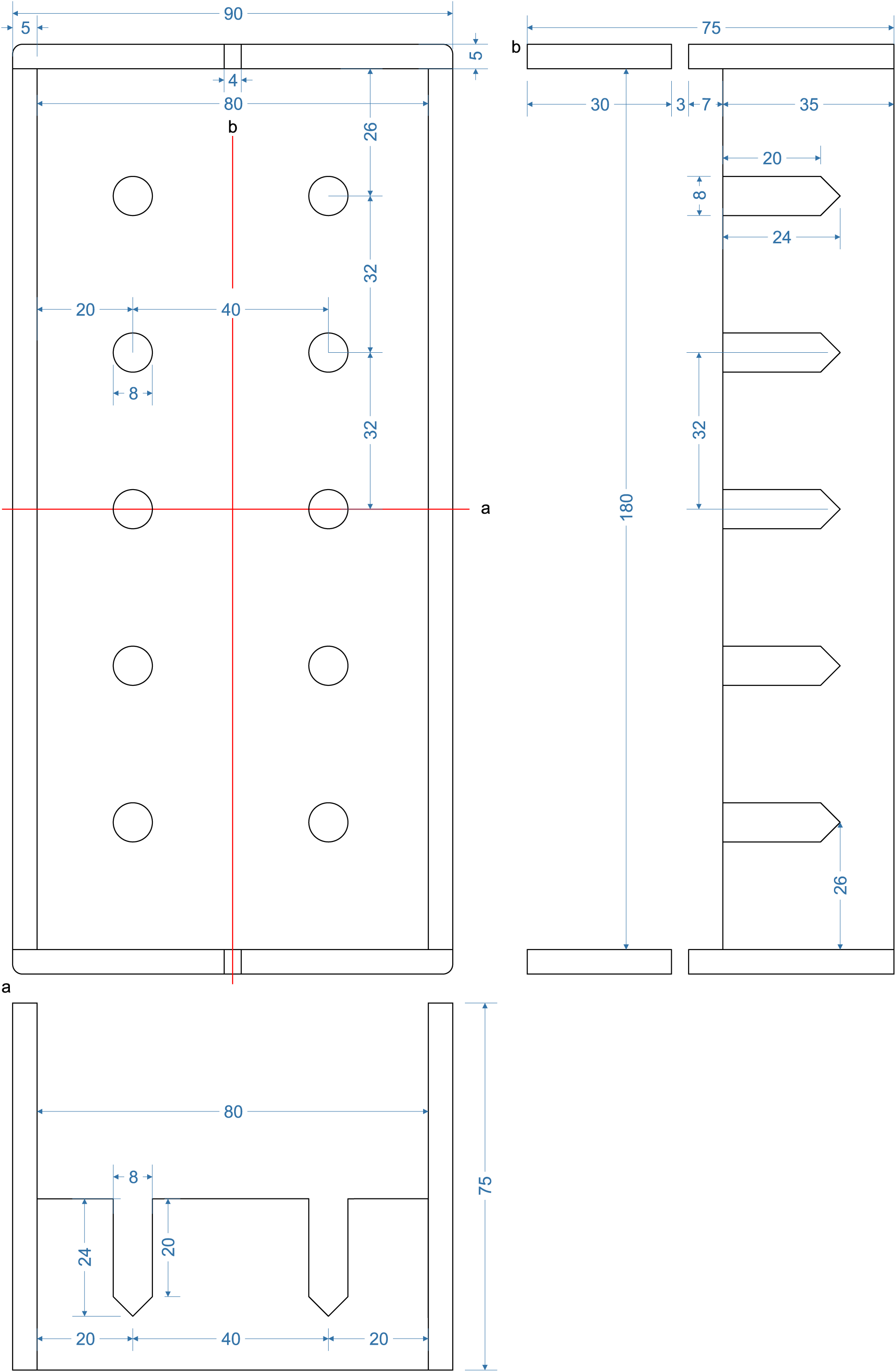

**Bottom part of pipette filling station.**

1,5 ml microcentrifuge tubes with intracellular solution are inserted into the 8 mm holes. Slight adjustment might be necessary to guarantee upright position of the tubes and easy insertion.

Distance between the holes (32 mm) are determined by the distance of the central channel between two connected Braun 3-way stopcocks. Slight variation might be necessary when different models are chosen. Height of the walls are determined by the length of the pipettes to ensure that pipettes will reliably dip into the tubes. The current measurements are optimized for approximately 40 mm long pipettes.

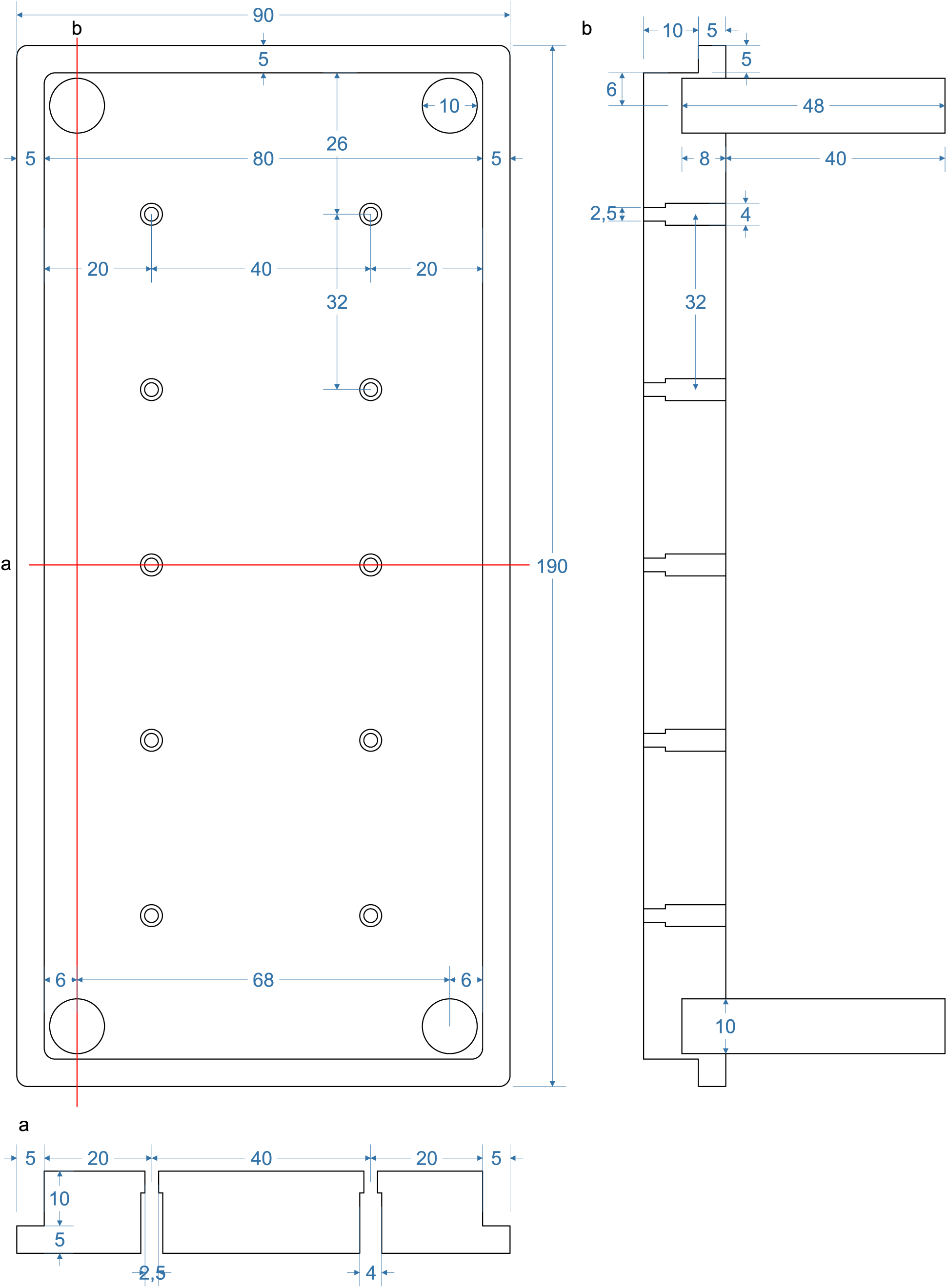

**Top part of pipette filling station (“lid”).**

Pipettes will be inserted into the 2,5 mm holes. Silicon tubes with 1,5 mm inner diameter and 3,5 mm outer diameter will be inserted into the 4 mm holes until the narrowing.

Section (a) depicts the horizontal section through the middle depicting the 2 mm and 4 mm holes.

Section (b) depicts a longitudinal section through the pillars of the lid. These enable a stable upside-down positioning of the lid in which the pipettes will be pointing upwards. The pillars are glued into a 10 mm hole.

#### Analog output routing device

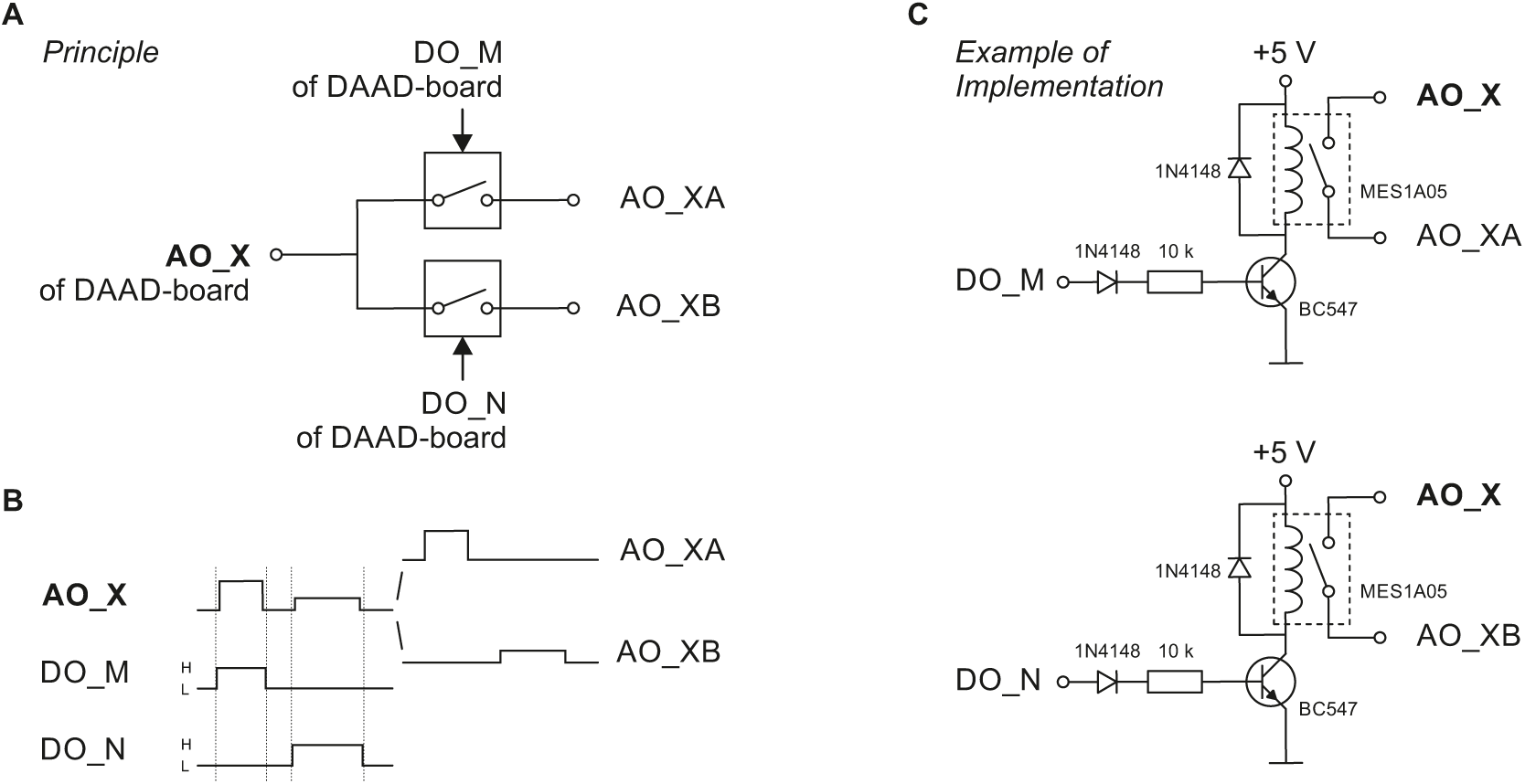

This device is necessary when more headstages are to be used in a multipatch setup than analog output channels are available on the digital-to-analog/analog-to-digital (DAAD-) board.

**A** Principle of a very simple and limited analog-out-switch distributing the analog out signal of the DAAD’s analog output channel AO_X to two channels serving two amplifiers or amplifier channels. The switches are gated by digital output channels of the DAAD-board (DO_M and DO_N).

**B** Via AO_X two different, non-simultaneous, non-overlapping command signals (different time of occurence, amplitude, duration) are sent to the switch, DO_M and DO_N are set high (H) as indicated (note that the digital pulses should start slightly earlier and terminate slightly later than the intended analog signals) and thus gate the respective command to the respective output of the switch (AO_XA or AO_XB). This switch is limited in the sense that the connected AO_XA and AO_XB will carry an identical (voltage- or current-) command signal (technically in both cases a voltage signal) from the DAAD to the individual amplifier inputs if command signals are to be applied simultaneously.

**C** Example of implementing the principle shown in A. To distribute the analog out command signal of the DAAD-board to two amplifier command input channels, two reed relais (e.g. MES1A05) are used. The power supply can be taken from an USB-port of a computer.

